# Major waves of H2A.Z incorporation during mouse oogenesis

**DOI:** 10.1101/2025.06.14.659461

**Authors:** Madeleine Fosslie, Erkut Ilaslan, Trine Skuland, Adeel Manaf, Mirra Søegaard, Marie Indahl, Maria Vera-Rodriguez, Rajikala Suganthan, Ingunn Jermstad, Shaista Khan, Knut Tomas Dalen, Ragnhild Eskeland, Michel Choudalakis, Magnar Bjørås, Peter Fedorcsak, Gareth D. Greggains, Mika Zagrobelny, John Arne Dahl, Mads Lerdrup

## Abstract

Mammalian oocytes and embryos have distinct epigenomes that undergo major transitions essential for transmitting life to the next generation. Here, we provide the first genome-wide maps of the histone variant H2A.Z during six different stages of mouse oogenesis. Utilizing picogram- scale chromatin immunoprecipitation and sequencing (picoChIP-seq), we reveal major waves of H2A.Z incorporation occurring early in growing oocytes. We identify distinct patterns of H2A.Z signal at oocyte-specific, embryo-specific, and constant H2A.Z loci. Late oocyte-specific loci precede reduced formation of lamina associated domains and early replication timing in the maternal compared to the paternal genome of 2-cell embryos. While constant H2A.Z is strongly associated with CpG islands (CGIs) and H3K4me3 near transcription start sites (TSS) of active genes, tens of thousands of oocyte- and embryo-specific H2A.Z incorporation sites exist independently of CGIs and TSSs. These TSS-distal H2A.Z sites are frequently enriched at transposable elements (TEs), and an intriguing inverse relationship exists between H2A.Z and H3K4me3 in low CpG environments, such as MTA and MTB retrotransposons (RTs). The existence of changes in H2A.Z distribution that persist across related developmental stages enable preservation of epigenetic information despite major concurrent changes in H3K4me3, H3K27me3, and DNA methylation. Altogether, this reveals new layers of regulation advancing our understanding of how histone variants contribute to the epigenetic landscape during mammalian oogenesis and preimplantation embryo development.

## Introduction

Mammalian oocytes and preimplantation embryos undergo extensive epigenetic changes, involving the establishment and remodeling of specific epigenetic marks crucial for development and fertility, reviewed in^1^. The histone variant H2A.Z is essential for embryo development, as H2A.Z knockout (KO) mouse embryos fail to develop beyond embryonic day 6.5^2^. H2A.Z has approximately 60% amino acid similarity to core histone H2A and is deposited in a cell-cycle independent manner, reviewed in^3^. H2A.Z can alter the structure and stability of chromatin and nucleosomes, recruits different types of reader proteins, and provides an obstacle for transcriptional elongation of the RNA pol II complex, reviewed in^4^. Accordingly, H2A.Z is involved in diverse biological processes, such as gene activation, firing of replication origins, DNA repair, meiotic recombination, embryonic stem cell differentiation, nucleosome turn-over, memory formation, chromosome segregation, heterochromatin silencing, and progression through the cell cycle^3, 5, 6^. H2A.Z exists as two isoforms in mice (H2A.Z.1 and H2A.Z.2), which are products of two nonallelic genes (*H2afz* and *H2Afv*) that encode proteins differing by only three amino acids and are differentially expressed across tissues^7–9^. In addition, the function of H2A.Z is affected by post-translational modifications and nucleosome partners^3^.

H2A.Z is generally present at promoters with CpG islands (CGIs) and at transcription start sites (TSSs) of active and poised genes in various cell types, including stem cells. In mammals, it usually colocalizes with the histone modifications H3K4me3 and H3K27me3^10–12^. H2A.Z can affect the deposition of H3K4me3 and H3K27me3, as it can facilitate recruitment of MLL and PRC2 complex proteins, which are responsible for the formation of H3K4me3 and H3K27me3, respectively^13^. The H2A.Z protein was shown to be present in MII oocytes^9^, but a recent genome-wide study did not detect genomic H2A.Z enrichment by ULI-NChIP-seq in MII oocytes despite a strong immunostaining signal^14^. Additionally, knockdown of H2A.Z at the zygote stage significantly reduces blastocyst formation, but has no effect on morula development, leading to the hypothesis that H2A.Z is not essential for initiation of zygotic genome activation (ZGA) in mice^14^. However, since the genome-wide dynamics of H2A.Z has not yet been analyzed in detail in developing mammalian oocytes, much remains to be learned about the possible roles at these stages.

Concurrent with the epigenetic changes in developing oocytes and embryos, transposable elements (TEs) are expressed in stage-specific waves, indicating precise regulation^15^. TEs are mobile multi copy DNA sequences that account for approximately half of the mammalian genome and are generally silenced to prevent indiscriminate activation and mutations by transposition, reviewed in^16^. Nonetheless, TEs are thought to promote evolution and facilitate organismal development by contributing enhancer and promoter sequences, modifying three-dimensional (3D) chromatin architecture, and giving rise to novel regulatory genes^16^. Among TEs, retrotransposons (RTs) constitute up to 38% of the transcriptome in mammalian preimplantation embryos^17^; indeed early development is marked by expression of the RTs ERVL, MaLR, and LINE1, some of which are fundamental for the progression of mammalian embryonic development, reviewed in^18, 19^.

During mouse oogenesis, primordial follicles assemble shortly after birth and develop into primary follicles. As a cohort of these follicles are activated, the oocyte within each begins to grow and increases its transcription during the subsequent postnatal (P) period, reviewed in^20^. The growing follicles reach the antral stage when a fluid-filled cavity forms. Inside the oocyte nucleus, known as the germinal vesicle (GV), chromatin condenses around the nucleolus. This transition marks a shift in chromatin organization from a non-surrounded nucleolus (NSN) to a surrounded nucleolus (SN) configuration. Concurrently, global transcription is silenced, and the oocyte reaches its full size around this time. In preovulatory follicles, oocytes resume meiosis in response to hormonal signals and progress to metaphase II (MII), where they are considered mature and remain arrested until ovulation and fertilization occur^20^. Nuclear organization is established *de novo* shortly after fertilization, reflected in the formation of Lamina Associated Domains (LADs) where the genome interacts with the nuclear lamina^21^. The fertilized oocyte relies on maternal RNAs until transcription is initiated during minor and major zygotic genome activation (ZGA) at the zygote and 2-cell stages, respectively, reviewed in^22^.

Given the relative scarcity of mouse oocytes and the limited sensitivity of chromatin immunoprecipitation and sequencing (ChIP-seq) assays, the genomic localization of H2A.Z and its relationship to the epigenome and transcriptional regulation in oocytes has remained unstudied. Here, we carry out picoChIP-seq to provide genome-wide maps of H2A.Z throughout mouse oogenesis by analyzing six oocyte stages, spanning from just after the oocytes have entered the growth phase until mature MII oocytes, as well as preimplantation embryo development. In contrast to the distinctive redistribution of many histone marks^23^, H2A.Z retains many canonical features during early development, including narrow peaks at TSSs and CGIs. Additionally, our maps reveal dynamic and stage-specific changes in H2A.Z deposition during oogenesis and early embryonic development. Interestingly, oocyte-specific H2A.Z is incorporated into regions in the genome where the embryonic formation of LADs differs markedly between the maternal and paternal genomes. Moreover, these loci replicate earlier in the maternal than the paternal genome, indicating that maternal H2A.Z incorporation or associated features may be instructive for the replication timing in the embryo. Much of the stage-specific H2A.Z incorporation occurs at non-TSS and non-CGI loci, which often correspond to TEs involved in regulation of transcripts crucial for oocyte transcriptome and epigenome establishment. Finally, we relate H2A.Z incorporation to genetic and epigenetic features and find an intricate relationship between H2A.Z, CpG density and H3K4me3. By characterizing these dynamic changes in H2A.Z distribution, this work highlights how histone variants contribute to the dynamic epigenetic landscape and genome regulation during oogenesis and preimplantation embryo development.

## Results

### picoChIP-seq reveals major waves of H2A.Z incorporation in growing oocytes

To assess the genome-wide profile of H2A.Z in limited numbers of mouse oocytes, we utilized our recently developed picoChIP-seq method to target H2A.Z^24^. We further validated the efficiency of picoChIP-seq for H2A.Z by successfully mapping H2A.Z in mouse ES cells chromatin from 1,000 and 500 cells, showing a comparable signal to that of published bulk H2A.Z ChIP-seq^13, 25^ (Fig. S1a,b,c). Next, we mapped H2A.Z in oocytes from mice at postnatal day 7 (P7), P10 and P12, and in NSN, SN, and MII oocytes, as well as in 8-cell, morula and blastocyst preimplantation embryos (Fig. 1a). This enabled us to resolve the oocyte GV stage and investigate whether the transition from the actively transcribing NSN -stage to the silenced SN -stage^20^ brings about any changes in H2A.Z localization. As H2A.Z ChIP-seq enrichment profiles have been characterized in mouse embryos, but not oocytes, we carried out cross-validation of our blastocyst H2A.Z picoChIP-seq data with recently published blastocyst H2A.Z ChIP-seq data^14^ (Fig. S1d,e). Our blastocyst H2A.Z ChIP-seq signal correlated well with that of published data, both when assessing the signal at TSSs (Spearman’s ρ = 0.92, ρ = 0.94) and throughout the entire genome (ρ = 0.82, ρ = 0.85, Fig. S1d,e).

**Fig. 1.**
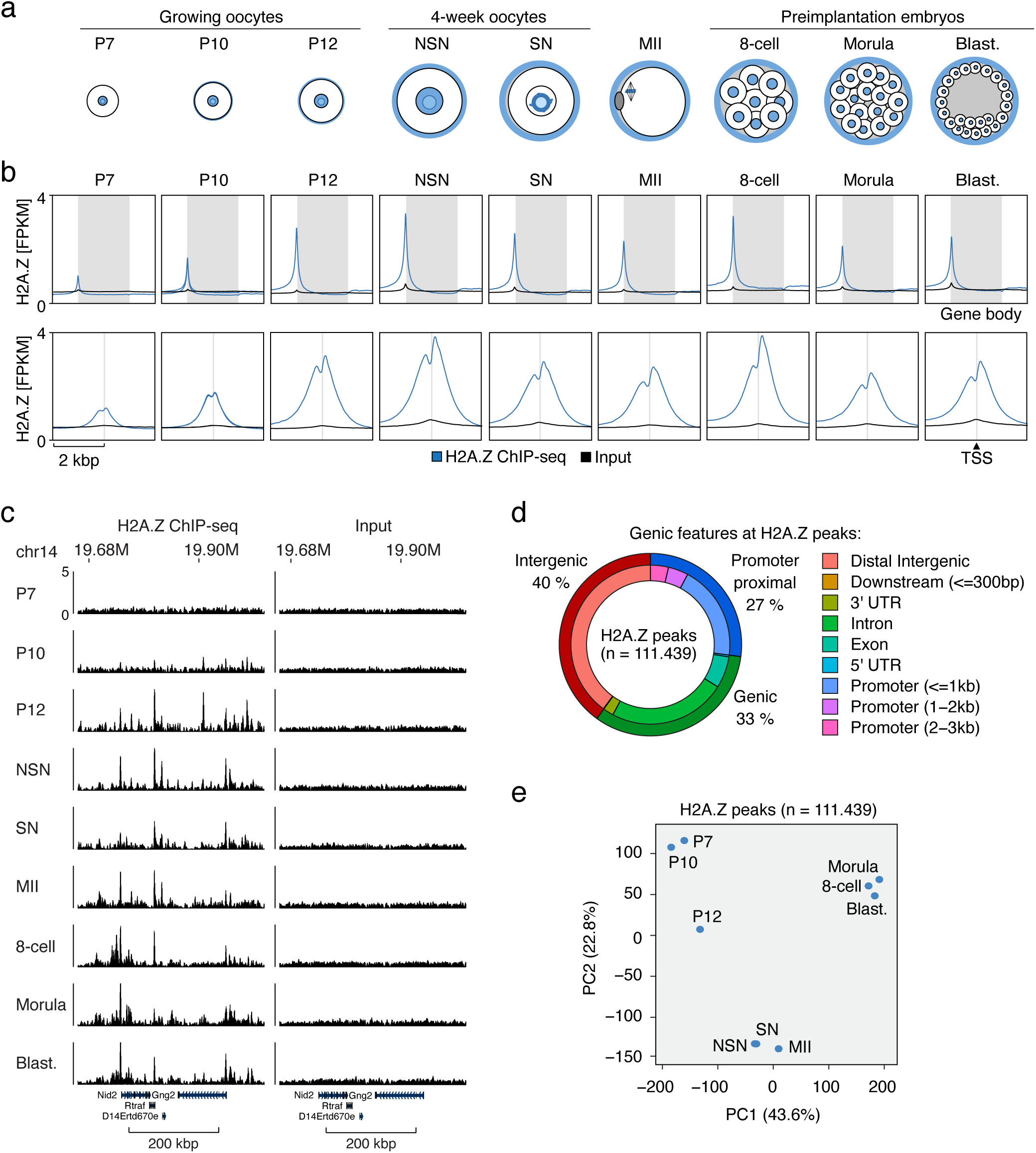
H2A.Z is incorporated both in mouse oocytes and preimplantation embryos. **a**, Overview of the developmental stages that were sampled and analyzed in this study. P, postnatal day; NSN, non-surrounded nucleolus; SN, surrounded nucleolus; MII, metaphase II; Blast., blastocyst. **b**, H2A.Z ChIP-seq signal in different developmental stages at gene bodies (shaded area, upper panel), n=47,248 and unique TSS +/-2kbp (shaded area, lower panel), n=32,166. **c**, Genome tracks of H2A.Z ChIP-seq signal from different developmental stages and at corresponding input samples, see also Fig. S1a. **d**, H2A.Z ChIP-seq peak overlap with genomic features in the mm10 genome, see also Figure S1f. **e**, Principal component analysis of the intensity of H2A.Z enrichment at H2A.Z peaks at different stages.

To investigate the location of H2A.Z in the mouse mm10 genome, we visualized the signal at gene bodies and unique transcription start sites (TSSs) (Fig. 1b). This was clearly enriched at TSSs (Fig. 1b) and varied during different developmental stages (Fig. 1b,c). Of note, we observed a major wave of H2A.Z establishment in growing oocytes, with minimal presence in P7, rapidly increasing to peaking levels in P12 and NSN oocytes (Fig. 1b). This reveals a profound switch in the incorporation of a histone variant in mammalian oocytes.

To identify regions in the genome with high H2A.Z enrichment, we performed peak calling for each oocyte and embryo stage and generated a combined peak set (n=111,439, Table S1) that included all peaks from all the developmental stages. In contrast to prominent histone modifications, H2A.Z incorporation generally occurred in relatively narrow regions with a mean size of 877 bp. Annotation of these peaks showed that H2A.Z frequently was incorporated away from genes and TSSs (Fig. 1d). Specifically, only 27% of H2A.Z peaks were located near promoters, while 33% and 40% were intragenic and intergenic, respectively (Fig. 1d). Principal component analysis (PCA) of the signal at these peaks revealed considerable stage-specificity in the H2A.Z localization (Fig 1e). Embryos at the 8-cell, morula and blastocyst stages showed a high degree of similarity but were distinct from that of all oocyte stages. Likewise, oocytes at later stages of oogenesis (NSN, SN, MII) clustered together, demonstrating a high degree of similarity. Growing oocytes (P7, P10, and P12), however, were clearly distinct from NSN, SN, and MII oocytes and relatively more heterogeneous in their H2A.Z localization. As growing oocytes are only separated by two to five days of growth, this supports a striking, rapid change in H2A.Z establishment early after oocytes enter the growth phase (Fig. 1e).

To assess the chromatin context in which the oocyte and embryo H2A.Z signal exists, we made use of the mouse full-stack ChromHMM annotation^26^. This revealed enrichment of H2A.Z at regions characterized as TSSs, promoters, enhancers and open chromatin, and H2A.Z depletion in regions characterized as heterochromatin (Fig. S1f). However, as these genomic states were defined using ChIP-seq data from somatic cells, the results mainly characterize the relationship to epigenomes of later developmental stages or adult tissues^1^.

### Oocyte and early embryo H2A.Z is incorporated at CGIs near TSSs of expressed genes

To visualize H2A.Z enrichment at TSSs over time, we performed k-means clustering of the TSS-associated H2A.Z signal across developmental stages using TSSs defined from annotated genes in the mouse mm10 genome (Fig. 2a). Surprisingly, few clusters showed predominant enrichment in either the developing oocyte (cluster VII) or the early embryo (clusters V and VIII), and many clusters had persistent enrichment throughout all stages (clusters I-IV and VI). Approximately 40% of TSSs had negligible H2A.Z signal and these TSSs were also lacking CGIs (clusters IX and X) (Fig. 2a). Next, we related the clustered TSSs to previously published RNA expression levels encoded by nearby loci^27^ and found an overall positive relationship between H2A.Z and RNA levels (Fig. 2b). Only one cluster (IV) displayed changing RNA levels, while the others were associated with constant RNA levels across the developmental stages. An investigation of developmentally important gene types within the clusters showed no specific enrichment or depletion in the oocyte or early embryo clusters. However, clusters II and III with constant RNA expression were depleted of maternal-specific genes and enriched in genes transcribed during major ZGA and mid-preimplantation gene activation (MGA), while clusters IX and X with low H2A.Z enrichment and low gene expression displayed the opposite pattern (Fig. 2c). Minor ZGA genes were exclusively enriched in cluster IV, with corresponding high RNA levels at the oocyte and zygote stage and low RNA levels at later stages. Overall, H2A.Z was located at genes expressed at ZGA and MGA both during oogenesis and preimplantation embryo development. Since we detected H2A.Z before the mouse minor and major ZGA, which occur in the zygote and the 2-cell embryo, respectively^22^, H2A.Z may contribute to marking these genes, as well as others, for later transcription, as reported in zebrafish and *Drosophila* embryos^28–30^. Notably, the relatively constant H2A.Z levels observed at many TSSs during oogenesis and preimplantation development contrast with the highly dynamic levels of epigenetic marks at these stages^23^. This stability may reflect a role for H2A.Z as a placeholder during epigenetic restructuring, as previously suggested in zebrafish^4^.

**Fig. 2.**
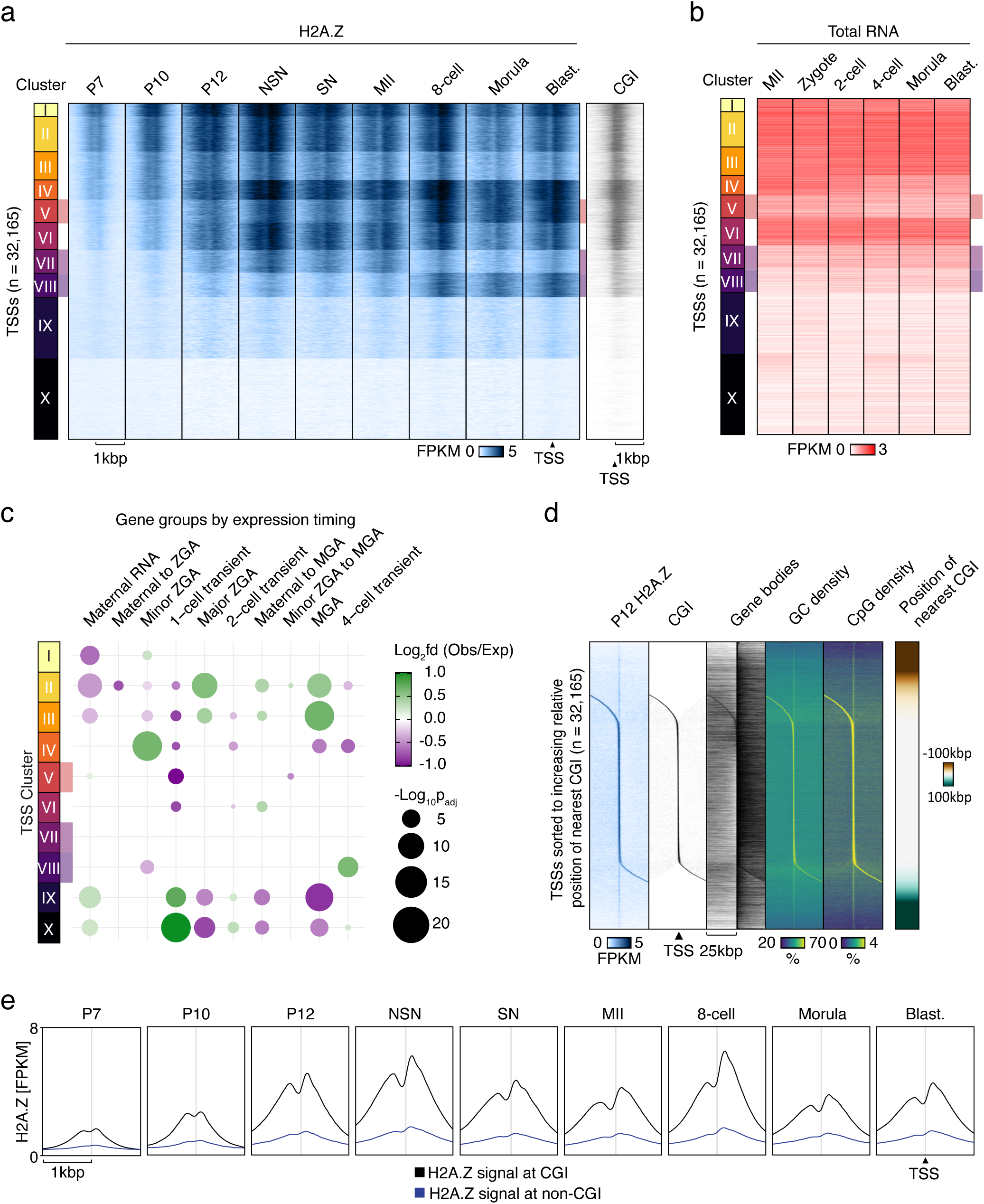
H2A.Z incorporation at TSSs of developmental genes and CGIs. **a**, H2A.Z ChIP-seq signal in different developmental stages at unique TSSs +/-1kbp (n=32,166), clustered based on H2A.Z signal. **b**, RNA levels^27^ at unique TSSs +/-10kbp (n=32,166), clusters based on Fig. 2a. **c**, Bubble plots showing overlap between TSS clusters and oocyte and embryonic gene categories compared to an average distribution across all clusters. p-values are calculated by Chi Square tests Benjamini-Hochberg adjusted for multiple testing. ZGA, zygotic genome activation; MGA, mid-preimplantation gene activation. **d**, Heatmaps showing H2A.Z, CGIs, Gene bodies, G/C, and CpG densities at unique TSS +/-25kbp (n=32,166) sorted according to increasing relative position of the nearest CGI. **e**, H2A.Z in different developmental stages at unique TSSs (n=32,166), either overlapping (black) or not overlapping (blue) CGIs.

To determine whether H2A.Z enrichment was specifically associated with TSSs, or alternatively with CGIs at or proximal to TSSs, we calculated the proximity of TSSs to CGIs and sorted them accordingly. It was clear that the majority of the H2A.Z signal was located on CGIs (Fig. 2d,e, S2a), whereas low-level enrichment was observed at TSSs that were not associated with CGIs (faint vertical line in Fig. 2d and S2a and blue lines in Fig. 2e). When visualizing CGIs, gene bodies, GC density and CpG density (Fig. 2d), we found that most of the H2A.Z signal at TSSs not associated with CGIs was instead enriched in CpG-dense regions that were not part of annotated CGIs. Moreover, a positive correlation was found between CpG density and H2A.Z enrichment at TSSs for P12 onwards (Spearman’s ρ=0.75, Fig. S2b), while the correlation was negligible in genome-wide assessments (Fig. S2c). An analysis of the standard deviation of the H2A.Z signal across the investigated oocyte and embryo stages showed that most of the variation in H2A.Z localization between stages takes place at non-TSSs loci (Fig. S2d). In summary, H2A.Z at TSSs was strongly associated with CGIs, and to a lesser extent, CpG-dense regions, while in other parts of the genome, H2A.Z exhibits a more variable distribution that was considerably less linked to CpG density.

### Dynamic H2A.Z incorporation reflects epigenetic changes and marks early replication and LAD formation in the embryo

Since CGIs near TSSs can only explain a minority of the H2A.Z localization (Fig. 1d), and the most variable signal exists elsewhere in the genome (Fig. S2d), we did an unbiased analysis of the full set of H2A.Z peaks from all oocyte and embryo stages. To this end, we performed k-means clustering of the H2A.Z signal at all H2A.Z peaks across all stages (Fig. 3a). This analysis revealed a clear pattern of stage-specific differences, with some clusters of H2A.Z enrichment being specific to early growing oocytes (cluster 1 and 2), further progressed and fully mature oocytes (cluster 3), as well as embryonic stages (cluster 8-10). Additionally, four clusters displayed a more constant H2A.Z signal from P12 onwards (cluster 4-7), and these clusters were strongly enriched for CpGs in comparison to the genome-wide mean (Fig. 3a). Related developmental stages showed high levels of similarity, as also observed in the PCA (Fig. 1e), demonstrating consistency in our data. As a control measure we also assessed the pattern of the H2A.Z signal in the two replicates of P10 oocytes from this study and found it to be highly similar (Fig. S3a). Likewise, H2A.Z enrichment patterns were similar between the blastocyst sample pool derived from this study and previous data^14^ (Fig. S3b). Altogether, these findings support the specificity of H2A.Z enrichment observed across developmental stages.

**Fig. 3.**
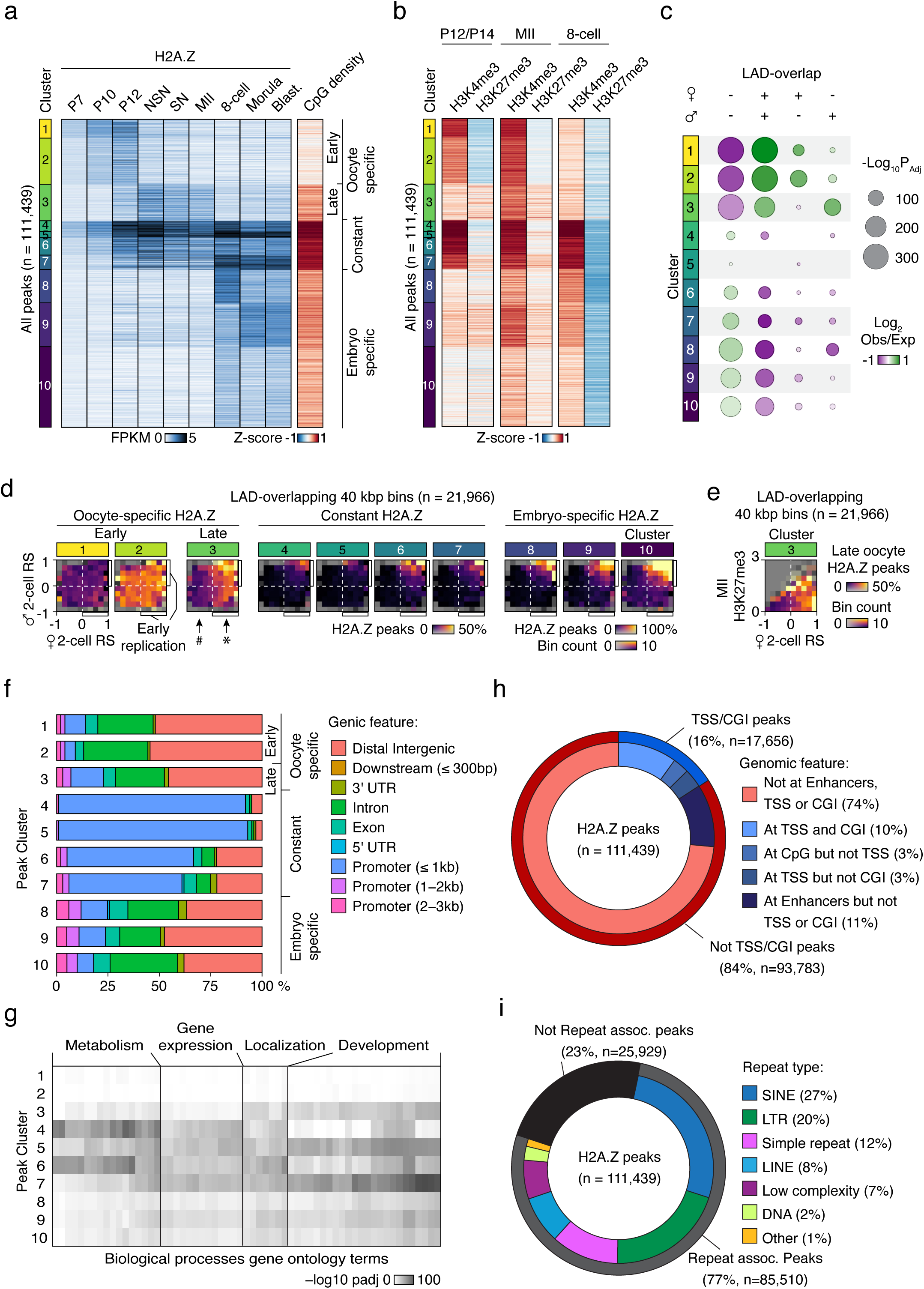
Oocyte and embryo specific H2A.Z peaks are mainly found outside of genes. Heatmaps of H2A.Z enrichment at an aggregated and clustered H2A.Z peak set from the indicated stages of oocyte and embryo development. **a**, Heatmaps of H2A.Z enrichment at aggregated and clustered H2A.Z peaks during different developmental stages as well as Z-score of CpG density at peaks. **b**, Z-scores of histone marks^31, 33, 54^ at clustered peaks. **c**, Bubble plots showing overlap of LADs and clusters of H2A.Z peaks compared to an average distribution across all clusters. p-values are calculated by Chi Square tests Benjamini-Hochberg adjusted for multiple testing^21^. **d**, 2D-histograms showing the genome-wide occurrence of each cluster of H2A.Z peaks (color) in relation to the mean Replication Status of the maternal (X-axis) and paternal (Y-axis) genomes^34^ in individual 2-cell embryos. # and * indicates noteworthy sex-specific differences, where low and high peak-occurrences, respectively, largely follows the maternal Replication Status, but not the paternal. Data were analyzed in 40 kbp bins, and the subset of bins overlapping with LADs are shown (n=21,966). For the non-overlapping subset see Fig S3d. **e**, 2D-histogram showing the genome-wide occurrence of cluster 3 H2A.Z peaks (color) in relation to the mean maternal Replication Status (X-axis) in individual 2-cell embryos and H3K27me3 levels in MII oocytes (Y- axis). Data were analyzed in 40 kbp bins, and the subset of bins overlapping with LADs are shown (n=21,966). For the non-overlapping subset see Fig S3d. **f**, Overlap between peaks in each cluster and indicated genic features. **g**, Heatmap of gene ontology (GO) analysis for H2A.Z peak annotated genes. –log_10_ transformed adjusted p-values of gene ontology biological process terms (X-axis) are visualized across H2A.Z peak clusters (Y-axis). Biological processes with a cumulative –log_10_ adjusted p-value in all clusters greater than 150 are visualized and labeled as general biological processes for clarity. **h**, Overview of the fraction of peaks that overlap with genomic features. **i**, Overview of the fraction of peaks that overlap with major classes of repeats.

In most cell types, the H3K4me3 histone modification is associated with active transcription, while H3K27me3 is associated with repressive chromatin and transcriptional silencing, and both marks are localized at CGIs^1^. During oogenesis and preimplantation embryo development, however, the distribution of H3K4me3^31, 32^ and H3K27me3^33^ becomes atypical and characterized by the presence of broad domains. To study incorporation of H2A.Z in the context of these extensive epigenetic changes, we assessed association between clusters of H2A.Z and previously published profiling of H3K4me3^31^ and H3K27me3^33^ in mouse oocytes and embryos (Fig. 3b). The dynamic H2A.Z signal was mirrored by changes in the epigenetic context, and we observed a strong stage-specific enrichment of H3K4me3 at H2A.Z peaks, while H3K27me3 was predominantly depleted, particularly in 8-cell embryos (Fig. 3a,b). Notably, early oocyte-specific H2A.Z peaks (clusters 1 and 2) displayed high to intermediate enrichment of H3K4me3 in P12 compared to the mean genome-wide signal, while P14 H3K27me3 was depleted at these peaks (Fig. 3b). The P12 pattern persisted to some degree to the MII stage, although both H3K4me3 and H3K27me3 exist as broad domains here. At the 8-cell stage, the H3K4me3 signal was reduced at early oocyte-specific peaks, while the H3K27me3 level was depleted in all H2A.Z clusters. The constant H2A.Z peaks in clusters 4-7 were characterized by strong enrichment of oocyte H3K4me3 as well as CpGs at all analyzed stages (Fig. 3a,b). Concordantly, clusters 4-7 were largely depleted when removing peaks overlapping with TSSs or CGIs from the peak set, (Fig. S3c). While H2A.Z signal was positively correlated to H3K4me3 in P12 oocytes and 8-cell embryos at TSSs (Spearman’s ρ = 0.83), correlations were reduced in the rest of the genome and at TSSs in MII oocytes (Fig. S3d). In conclusion, this suggests a closer relationship between H3K4me3 enrichment compared to H3K27me3 enrichment for the dynamic H2A.Z signal.

Nuclear organization is established *de novo* shortly after fertilization with the formation of (LADs), which are often related to gene repression^21^. Interestingly, the LAD formation of the two parental genomes differs until the 8-cell state^21^. To investigate if oocyte H2A.Z is associated with LAD formation, we analyzed the level of overlap between clustered H2A.Z peaks and LADs that were specific or shared between the two parental genomes in 2-cell embryos (Fig. 3c). We observed a highly significant co-occurrence between H2A.Z peaks arising in mature oocytes (cluster 3) and LADs that were specific for the paternal genome but not the maternal genome. As oocyte-specific H2A.Z peaks are maternally inherited, this indicates that factors in this genome and/or chromatin counteracted LAD formation (Fig. 3c). This was seemingly unrelated to the presence of broad H3K4me3 domains in the MII oocytes, since early oocyte H2A.Z peaks in cluster 1 and 2 underwent highly significant LAD-formation in the maternal genome only, despite being characterized by abundant MII H3K4me3 (Fig. 3c).

Recent investigations of replication timing in mouse early embryos revealed the emergence of a timing program in 2-cell embryos correlating with transcription, LADs, genome compartmentalization and inherited histone modifications with consistent differences between the parental genomes^34^. Given the involvement of H2A.Z in the activation of replication origins and replication timing^5^, we next investigated if H2A.Z played a role in replication timing. As LADs and late replication are strongly correlated^34^, we decided to analyze the parts of the genome overlapping and not overlapping with maternal 2-cell LADs separately (Fig. 3d, S3e). When relating the occurrence of the different developmental stage-specific H2A.Z peaks to the replication timing of the maternal and paternal genomes in the 2-cell embryo, we discovered a clear relationship between the maternal replication timing and the H2A.Z peaks occurring in mature oocytes (cluster 3) (Fig. 3d, maternal late and early replication indicated by # and *, respectively). In contrast, no such relationship was seen for early oocyte-specific H2A.Z peaks (clusters 1 and 2) or for the paternal genome (Fig. 3d). Also, peaks from clusters with more constant H2A.Z (clusters 4-7) or embryo-specific H2A.Z (clusters 8-10), were largely associated with the early replication of both the maternal and paternal genomes. The asynchronous early 2- cell replication of the maternal genome is defined by maternally inherited MII oocyte H3K27me3^34^, which recently was shown to antagonize LAD formation in the embryo^35^. Accordingly, we investigated whether the mature oocyte-specific H2A.Z incorporation was linked to asynchronous early maternal 2-cell replication through H3K27me3. While oocyte-specific H2A.Z peaks were frequent in such genomic regions characterized by H3K27me3 in MII oocytes, these peaks were also observed in counterparts with little or no preceding H3K27me3 in MII oocytes (Fig. 3e, S3f). Altogether, these intriguing relationships imply that maternally inherited H2A.Z incorporation or associated features may be instructive for the replication timing in the embryo through a mechanism independent of H3K27me3.

To characterize stage-specific and constant H2A.Z further, we analyzed the occurrence of genic features for each cluster. We found that the majority of the oocyte and embryo stage-specific H2A.Z incorporation took place at distal intergenic and intronic regions, while the constant clusters 4-7 were strongly enriched in promoters (Fig. 3f). To explore potential biological implications of H2A.Z incorporation, we next annotated peaks to TSSs of genes within 5kbp of the peaks and performed a gene ontology analysis for each peak cluster (Fig. 3g, Table S2). We found an overall enrichment of H2A.Z peaks at genes frequently involved in metabolism, gene expression, localization, and development-related biological processes. Intriguingly, the early oocyte-specific clusters (1 and 2) did not show any strong enrichment, while the late oocyte-specific cluster 3 showed an enrichment for localization and development-related genes, but limited association to biological processes related to gene expression. This analysis also revealed two different subsets of the more constant clusters, namely clusters 4 and 6 that contrast to cluster 5 and 7. While clusters 4 and 6 showed a strong enrichment in genes involved in metabolism related processes, clusters 5 and 7 were enriched in genes involved in development-related processes. Taken together, this analysis highlights that both stage-specific and constant presence of H2A.Z peaks is coupled to biological context of nearby genes involved in oogenesis and early embryonic development.

We next conducted a comprehensive analysis of genomic features at or near non-TSS/CGI H2A.Z peaks and confirmed the enrichment of H2A.Z at enhancers as well as in open chromatin regions (Fig. S3g,h). At enhancers^36^, the signals were highly dynamic, indicating involvement in gene regulation (Fig. S3g). The most notable changes at enhancers occurred between early growing oocytes, further progressed oocytes, and embryonic stages, while enrichment was highly similar within each of these groups. Moreover, non-TSS/CGI H2A.Z peaks were often associated with genes involved in transcriptional regulation and cell morphology (Fig. S3i). While 16% of H2A.Z peaks overlapped with TSS or CGIs and 11% with enhancers, 74% were located in other genomic regions (Fig. 3h). To identify possible features shared by the large fraction of H2A.Z peaks that did not overlap with TSSs and CGIs, we analyzed the overlap between this subset and repeat annotation from Repeatmasker^37^. We found that 77% of H2A.Z peaks colocalized with repeats, mainly SINE and LTR TEs (Fig. 3i). Only 18% of H2A.Z peaks did not colocalize with either a repeat, an enhancer, a TSS or a CGI. Thus, the majority of H2A.Z deposition in the genome of mouse oocytes and early embryos occurs at TEs and other repeat elements.

### Stage-specific H2A.Z at LTR retrotransposons

Given that TEs are expressed in stage-specific waves in mammalian embryos, with specific ones being essential for embryonic development^18^, we investigated which repeat types were enriched in our stage-specific H2A.Z peak clusters (Fig. 4a,b). The most abundant repeat types were long terminal repeat (LTR) RTs, particularly of the RMER and MT types. This was specific for H2A.Z established in early growing oocytes (clusters 1 and 2) (Fig. 4b, S4a). Thus, the major wave of H2A.Z establishment in growing oocytes is associated with RTs. The H2A.Z specifically incorporated during later oocyte stages (cluster 3), occurred in parts of the genome defined by simple repeats and low complexity regions (Fig 4b, S4a). To explore the underlying DNA sequences, we performed *de novo* motif enrichment analysis of clustered non-TSS/CGI peaks and found an increased frequency of GC rich motifs in peaks belonging to clusters 3, 8, 9 and 10 (Fig. S4b). However, a three-way analysis of the genome-wide relationship between H2A.Z, CpG densities, and GC% revealed that H2A.Z was largely associated with CpG density and not GC%, evident from the high GC% regions with low CpG density having lower H2A.Z signals than high CpG regions (Fig. S4c). These findings suggest that the association of H2A.Z to certain repeat types may be due to elevated CpG densities within repeats rather than specific motifs.

**Fig 4.**
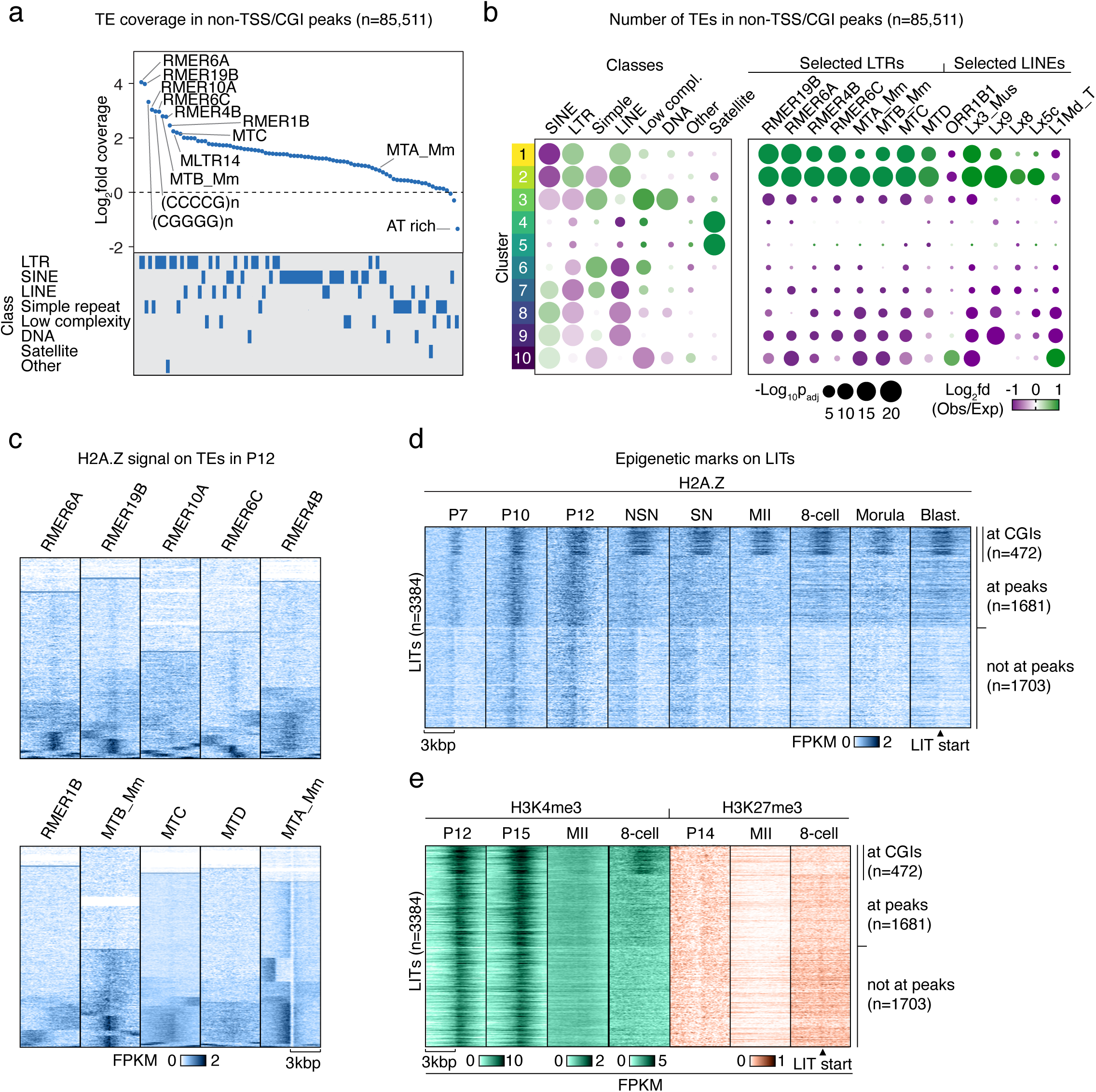
Oocyte-specific H2A.Z signal is enriched at specific repeats. **a**, Log_2_fold difference between observed and expected coverage of 90 repeat types with >200 counts in the non-TSS/CGI H2A.Z peak set. The bottom panel shows the repeat class. **b**, Bubble plots showing overlap of TE classes (left) and selected LTR and LINE subtypes (right) in clusters of non-TSS/CGI H2A.Z peaks compared to an average distribution across all clusters. p-values are calculated by Chi Square tests Benjamini-Hochberg adjusted for multiple testing. A complete plot of all TE types can be found in Fig. S4a. **c**, Heatmaps showing H2A.Z enrichment at selected RTs during the P12 stage clustered based on the H2A.Z profile at the start of the RTs +/-3kbp. **d**, Heatmaps of H2A.Z enrichment profiles at the start of LITs^39^ sub grouped based on overlap to CGIs and H2A.Z Peaks. **e**, H3K4me3^31^ and H3K27me3^33, 54^ profiles at LITs ordered as Fig. 4d.

The high copy number of TEs provided an ideal basis for systematically probing relationships between H2A.Z, histone marks, and transcription. We therefore investigated the localization and dynamics of H2A.Z in selected populations of enriched TE types and observed a localized enrichment near a considerable fraction of RMERs and MTs (Fig. 4b,c). Indeed, a large fraction of MTAs, which is the youngest and most abundant MT subfamily^38, 39^, were associated with dynamic H2A.Z enrichment across oocyte and embryo stages (Fig. 4c, S4d). H2A.Z levels differed in timing and localization relative to the MTAs (Fig. S4d), with the most pronounced and proximal enrichment occurring early in oocyte development and peaking at P12 stage (Fig. S4d, see *). Despite the many distinct H2A.Z profiles associated with MTAs, the relationship between H2A.Z profiles and transcription was limited, as many H2A.Z-negative MTAs were transcribed at the same level as H2A.Z-positive MTAs (Fig. S4d). In the P12 stage oocytes, increased upstream H2A.Z enrichment was associated with reduced upstream expression, suggesting that H2A.Z may strengthen transcriptional directionality at MTAs here (Fig. S4d, see *). These findings indicate an intriguing relationship between active RTs and H2A.Z establishment in growing oocytes, where the strongest H2A.Z enrichment is associated with absence of transcription.

Mammalian oocytes have a distinct transcriptome with an abundance of LTR-initiated transcription units (LITs), which are genic and intergenic transcripts initiating from solo LTRs^38,39^. Of the 3384 LITs previously identified in the mouse genome, many are MTAs (n=2301). When visualizing H2A.Z specifically at LITs, narrow and localized H2A.Z signal was generally evident at the P10 and P12 stages (Fig. 4d). Approximately half of these LITs (1703) did not overlap with detected H2A.Z peaks (Fig. 4d), but a low level of sub-threshold H2A.Z signal was still evident. Strong H3K4me3 enrichment in P12 and P15 oocytes within LITs showed that this epigenetic mark coexists with H2A.Z at a considerable fraction of these elements (Fig 4e). Likewise, the LITs with H2A.Z signal at CGIs (472) (Fig. 4d) displayed both H2A.Z and H3K4me3 enrichment during early oogenesis and embryo development, only interrupted at the MII stage by the formation of broad H3K4me3 domains. In contrast, H3K27me3 signal remained low within LITs (Fig. 4e). These findings show a dynamic pattern of H2A.Z and H3K4me3 at LITs with a considerable degree of colocalization.

### H3K4me3-independent H2A.Z is incorporated into low-CpG loci at MTAs and MTBs

In contrast to TSSs, where we found a strong correlation between H2A.Z and H3K4me3, the strength of the association between H2A.Z and H3K4me3 was reduced at the genome-wide level (Fig. S3d). To investigate this in greater detail, we related genome-wide H2A.Z and H3K4me3 signal in P12, MII and 8-cell stages to CpG enrichment (Fig. 5a). Interestingly, the parts of the genome with the lowest levels of H2A.Z and H3K4me3 coexistence were also characterized by low CpG density in P12, MII and 8-cell stages, whereas coexistence generally was associated with high CpG densities (Fig. 5a). To probe these intriguing relationships further, we again took advantage of the high copy numbers of RTs and clustered selected RTs based on the combined profiles of H2A.Z, H3K4me3 and H3K27me3. Visualization of common epigenetic marks at the clustered RTs in growing oocytes, revealed that regions with high levels of either H2A.Z or H3K4me3 were low in H3K27me3 and DNA methylation, and vice versa (Fig. 5b, S5). Furthermore, H2A.Z and H3K4me3 had minimal colocalization, especially at MTA RTs (Fig. 5b, S5). While both H2A.Z and H3K4me3 signals were generally present at the same loci, a notable spatial separation existed with H2A.Z mainly situated upstream of H3K4me3 (Fig 5b, c, clusters E, G and J). Moreover, H3K4me3 and RNA levels tended to be associated with an elevated abundance of CpG and GC at the regions flanking these MTA subgroups. H3K4me3 was largely absent in the approximately 30% of MTA loci displaying high H2A.Z signal (Fig. 5c, clusters F and I), and only the minor cluster H showed colocalization of H3K4me3 and H2A.Z. Increased RNA levels were mainly observed at loci in clusters E and G, generally at the parts enriched in H3K4me3, and a similar trend was observed for MTB (Fig. S5). When examining older types of RTs such as MTC and MTDs^38^, a greater fraction of the H2A.Z signal overlapped with H3K4me3 (data not shown), suggesting that lack of overlap between H2A.Z and H3K4me3 is a feature of younger types of MT RTs. In contrast, RMERs, Lx3_Mus and Lx9 LINE RTs demonstrated an increasing overlap of H2A.Z and H3K4me3 signals (Fig. 5b, S5), indicating a different regulatory mechanism for MTA and MTB RTs. In comparison, H2A.Z and H3K4me3 generally coexisted at CGIs (Fig. 5d), altogether demonstrating the existence of an intricate sequence-dependent relationship between the histone variant H2A.Z and the histone modification H3K4me3.

**Fig. 5.**
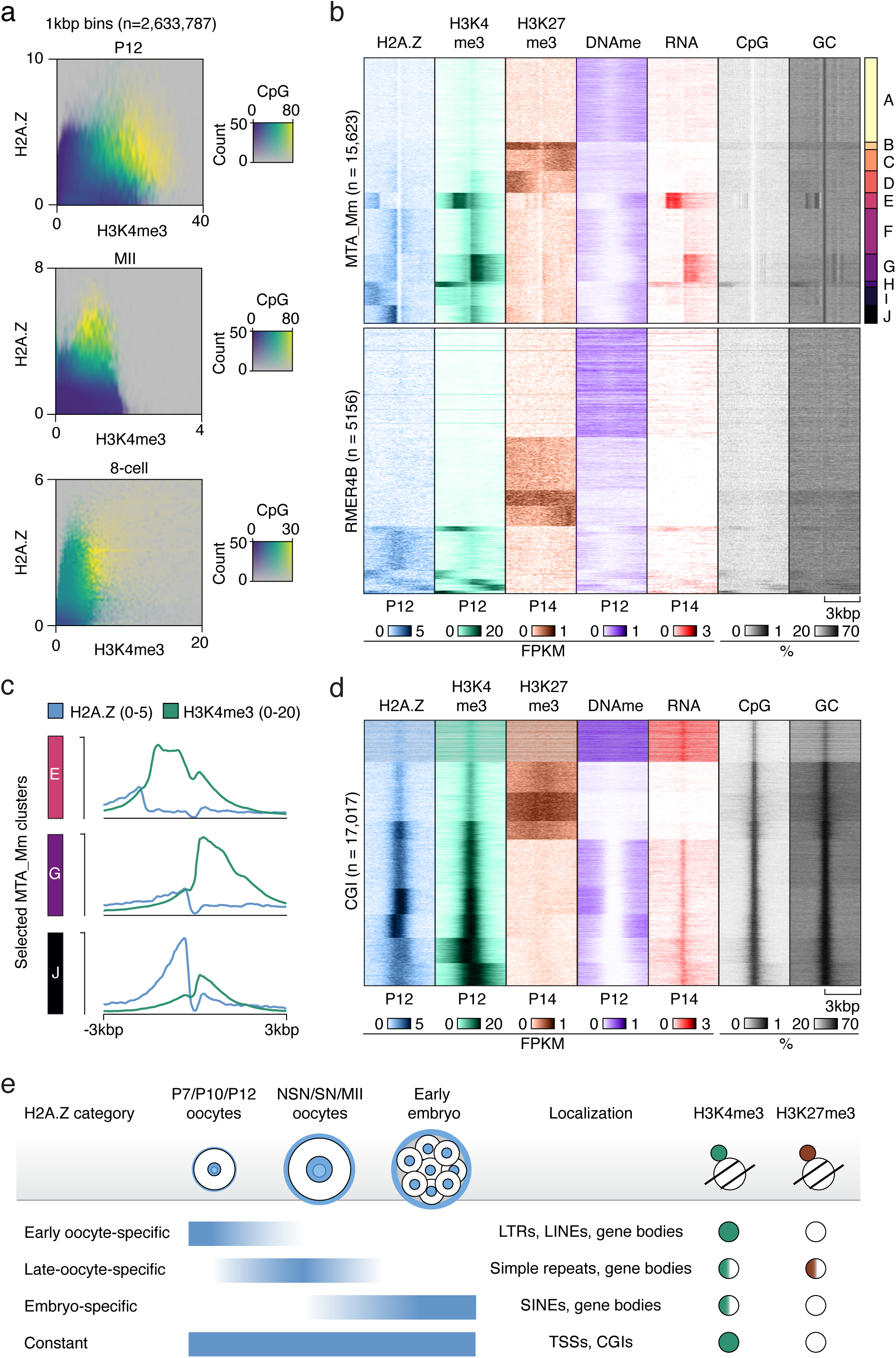
H2A.Z is not correlated to H3K4me3 and CpG density at many loci. **a**, Heatmaps showing the genome-wide relationships between H2A.Z, H3K4me3^31^ and CpG density in selected developmental stages visualized in 1kbp bins (n=2,633,787). Opacity reflect the occurrences of indicated H3K4me3 and H2A.Z levels. **b**, Heatmaps showing indicated features^27, 55^ at selected RTs clustered based on the combined distribution of H2A.Z, H3K4me3^31^ and H3K27me3^54^ at the start of each RT +/-3kbp in growing oocytes (P12 or P14). **c**, Average H2A.Z and H3K4me3 signal from selected clusters from 5b. **d**, Heatmaps showing indicated features at CGIs clustered based on the combined distribution of H2A.Z, H3K4me3^31^, and H3K27me3^54^ at the start of each CGI in growing oocytes (P12 or P14). **e**, Model of H2A.Z in mouse oocytes and the preimplantation embryo and its association with histone marks, CGIs, TSSs and TEs.

## Discussion

In this study, we provide the first genome-wide maps of the histone variant H2A.Z during six different stages of mouse oogenesis. Our work demonstrates a rapid major wave of establishment of H2A.Z early in oogenesis, peaking in P12 and NSN oocytes just prior to global transcriptional silencing. The landscape of H2A.Z signals is generally comprised by narrow peak profiles and enrichment at TSSs and CGIs. In this, H2A.Z differs markedly from previously assessed epigenetic marks at these developmental stages, including H3K4me3, H3K27me3, and DNA-methylation, which all display broader or otherwise non-canonical distributions that are unique for oocytes and preimplantation embryos^23^. Most importantly, this may allow epigenetic information to be preserved across these stages despite major concurrent changes in H3K4me3, H3K27me3, and DNA methylation. H2A.Z may play a crucial role in the regulatory dynamics, potentially acting as a placeholder that directs where future epigenetic marks should be deposited during the restructuring of the epigenome. Indeed, such a function has been proposed in zebrafish and *Drosophila* embryos, where H2A.Z is essential for ZGA and is detected at genes that will later become active or poised^28–30^. This function may be particularly important for the H2A.Z-enriched loci that characterize the minor ZGA, major ZGA, MGA, and the MT subfamily of RTs observed in our study, as these loci all become active in the zygote or shortly thereafter.

Our findings also reveal dynamic, stage-specific changes in H2A.Z profiles during oogenesis and preimplantation embryo development (Fig. 5e). Intriguingly, Oocyte-specific H2A.Z is strongly correlated to paternal LADs and early maternal replication timing, indicating that maternal H2A.Z incorporation or associated features may be instructive for the replication timing in the embryo. Additionally, these dynamic changes are particularly striking at non-TSS loci with low CpG enrichment, which often harbor TEs. There is growing evidence that RT TEs are intimately intertwined with mammalian embryonic development (reviewed in^40^). For example, the LINE family of RTs are transcriptionally active early in preimplantation development, peaking at the 2-cell stage in mouse^41^. Furthermore, H2A.Z is enriched at LINE L1 promoter regions in mouse cell lines, possibly associated with silencing of these young RTs^42^, while young RTs were also shown to be marked by H2A.Z in zebrafish embryos^43^. In our study, we identify several LINEs enriched for H2A.Z in the early oocyte-specific clusters (Fig. S4a), particularly Lx3_Mus and Lx9, with lower enrichment observed for Lx5c and Lx8. However, Lx RTs are older LINEs^44^, and we observed little accumulation of transcripts at these loci (Fig. S5). We also observe H2A.Z signal at RMERs and MTs (Fig. 4b), which are crucial for regulating the oocyte transcriptome and epigenome, and is frequently found in TE-initiated transcripts in mouse embryos^15, 38, 39^. The MT subfamily accounts for 13% of all transcripts in the fully grown oocyte^45^ and has been co-opted to drive expression during ZGA and early embryogenesis^46, 47^. The incorporation of H2A.Z at these RT TEs, important for embryonic development, further indicates that H2A.Z may be involved in the regulation of these critical developmental stages.

The non-TSS loci with H2A.Z deposition at RTs exhibited rather distinct and sometimes intriguing patterns of colocalization with H3K4me3: some followed a conventional pattern with H2A.Z and H3K4me3 overlap, some displayed H2A.Z positioned upstream of H3K4me3, and others lacked H3K4me3 entirely. While LITs were enriched in regions where H2A.Z overlapped with H3K4me3, an inverse relationship between H2A.Z and H3K4me3 was particularly clear in low CpG environments, such as at MTA and MTB RTs. The expressed MT RTs predominantly exhibited H3K4me3 downstream of H2A.Z (Fig. 5b,c, clusters E and G), except for a small subset displaying the conventional pattern (Fig. 5b, cluster H). These may correspond to the MTs previously shown to be active during early embryonic development^17^. Based on these observations, we propose a dynamic equilibrium model in which conventional H2A.Z is incorporated at CGIs that colocalize with H3K4me3 and active gene expression, but is continuously evicted by transcription and replenished through recruitment from CGIs. In contrast, H2A.Z not overlapping with H3K4me3 is found in GC-poor regions upstream of H3K4me3, likely due to H2A.Z eviction at TSSs by active transcription and reduced incorporation rates caused by low CpG density. In these regions, H2A.Z may regulate the transcription directionality.

Overall, the distinct patterns of H2A.Z observed at TSS-associated CGIs and CpG depleted TEs as well as correlating to LADs and maternal early replication timing, indicate that H2A.Z serves different functions across genomic regions and developmental stages. This may imply a role for H2A.Z in linking regulation of TEs to the broader epigenetic program, particularly during oogenesis, thereby laying the foundation for successful embryonic development. By characterizing these dynamic changes in H2A.Z distribution, this work advances our understanding of how histone variants contribute to the establishment and maintenance of epigenetic landscapes in oocytes and preimplantation embryos.

### Limitations of the study

Antibodies against H2A.Z do not distinguish between the two different, non-redundant isoforms of H2A.Z in mice. Furthermore, these isoforms may differ in their association with H3K4me3 and have been shown to be differentially expressed across tissues^3, 7, 48^. Accordingly, we cannot formally rule out the possibility that the observed differences in H2A.Z patterns are influenced by distinct functional roles of the two isoforms. Moreover, the antibody used in this study does not distinguish between different posttranslational modifications of H2A.Z, nor does it differentiate whether H2A.Z is incorporated into homotypic (H2A.Z/H2A.Z) or heterotypic (H2A/H2A.Z) nucleosomes, or whether it is combined with other canonical histones or their variants (e.g. H3.3). These combinations influence nucleosome stability, which in turn can impact the transcription of nearby genes^13, 49–51^.

## Methods

### Mouse Housing and Care

All mouse experiments were approved and registered by the Norwegian Food Safety Authority (NFSA approved application/FOTS IDs: 7216, 10898, and 24911, and 8743) and conducted in accordance with Norwegian regulation FOR-2015-06-18-761, which closely aligns with EU directive 2010/63/EU on the protection of animals used for scientific purposes. Housing for the C57BL/6NRj mice (Black 6N, Janvier Labs, France) and RjOrl:SWISS mice (CD-1^®^, Janvier Labs) was provided in individually ventilated cages (Green Line IVC SealSafe Plus Mouse, Tecniplast) connected to an Aero IVC air handling unit, under specific pathogen-free conditions. The cages were equipped with wood bedding material and enriched with plastic shelters, wooden sticks, and nesting paper. Standard housing conditions were maintained with 22 ± 2 °C room temperature and 55 ± 5 % relative humidity, on a light:dark cycle of 12:12 h light (7 AM:7 PM), and free access to water and rodent chow. The staff of the animal facility at the Department of Comparative Medicine, Oslo University Hospital and University of Oslo kept and cared for the mice.

### Collection of Mouse Oocytes

#### Strains and treatments

Postnatal day 7 (P7), P10, and P12 CD-1 females were used for collection of growing oocyte (GO) sample pools. Black 6N females aged 4 weeks were used for collecting NSN, SN and MII oocytes (Table 1). To induce superovulation for collection of these three stages, 5 international units (IU) of pregnant mare serum gonadotropin (PMSG; Prospec, HOR-272) were injected intraperitoneally (IP) at 2 pm. For NSN and SN stages, the oocytes were collected ∼45 hours after the PMSG injection. For collection of MII stage oocytes, an additional intraperitoneal injection with 5 IU of human chorionic gonadotropin (hCG; Sigma, C1063) was given 45-48 hours following the PMSG injection. Subsequently, MII oocytes were collected 18-22 hours after the hCG injection.

**Table 1.** Collected samples pools.

| Table 1. Collected samples pools |  |  |
| --- | --- | --- |
| Developmental stage | No. of cells | Mouse strain |
| P7 | 250 | CD1/Swiss |
| P10 | 250 | CD1/Swiss |
| P12 | 200 | CD1/Swiss |
| NSN | 200 | C57BL6/N |
| SN | 200 | C57BL6/N |
| MII | 300 | C57BL6/N |
| 8-cell | ~600 | C57BL6/N |
| Morula | ~110 | C57BL6/N |
| Blastocyst | ~460 | C57BL6/N |

#### Mouse GO collection

Dissected ovaries, 5 or 10 respectively, were placed in 3.5 mm dishes, and were enzymatically digested in 0.08 % Trypsin (Merck) at 37°C for 30 minutes while the ovaries were punctured with a 30G syringes (BD Micro-Fine+, Becton Dickinson). This was followed by 20 min treatment with 42 U/mL DNase (Sigma, 04716728001) at 37 °C (5% CO2), then a 10 min incubation with a final concentration of 1 mg/ml collagenase (Sigma, C9407) at 37°C (5% CO2) before collection, washing, and cross-linking as described below (picoChIP-seq > *Oocyte and embryo cross-linking*).

#### Mouse NSN/SN oocyte collection

Dissected ovaries with surrounding tissues were kept in M2 medium at 37°C prior to processing, with approximately four organs being processed at a time. With the aid of a heated stage stereomicroscope (37 °C), fat and surrounding tissues were removed carefully, and the ovaries were punctured with a 30G syringe (BD Micro-Fine+, Becton Dickinson) to mechanically release the oocytes into M2 medium supplemented with 0.2 mM 3-isobutyl-1- methylxanthine (IBMX; Sigma, l5879) to prevent meiotic resumption. Transfer to fresh, pre-warmed M2 medium with 0.2 mM IBMX was used for additional washes at 37 °C (no CO2). The presence or absence of a perivitelline space (PVS), defined as a visible gap between the oolemma and the zona pellucida, was used to sort cumulus-free oocytes into NSN or SN stage. PVS serves as a reliable approximation for NSN/SN staging, as the ability to form a PVS within 1 hour of in vitro culture with IBMX has been shown to correlate strongly with an SN chromatin configuration (Inoue et al., 2007). Finally, Tyrode’s acid (Sigma, T1788) was employed to remove the zona pellucida, followed by washes in fresh M2 and cross-linking using formaldehyde, as described below (picoChIP-seq > Oocyte and embryo cross-linking).

#### Mouse MII oocyte collection

In a petri dish with M2 medium (Sigma, M7167), the oviducts were transferred and examined using a heated stage stereomicroscope (37 °C) to identify the ampulla. Then, the oocytes were mechanically released, and the surrounding cumulus cells enzymatically removed using 0.3 mg/ml hyaluronidase (Sigma, C1063) in M2, incubated at room temperature. Transfer to fresh M2 medium was used for additional washes. Finally, Tyrode’s acid (Sigma, T1788) was employed to remove the zona pellucida, followed by M2 washes and cross-linking using formaldehyde, as described below (picoChIP-seq > *Oocyte and embryo cross-linking*).

### Mouse embryo culture and collection

Follicle growth was stimulated in 3.5 weeks old females (Black 6N, Janvier Labs) by an intraperitoneal injection with 5 IU PMSG at 2pm. Around 45 h later, 5 IU hCG (Chorulon, MSD Animal Health) was injected to induce ovulation. Directly after hCG injection, females were housed 1:1 with stud males of the same strain. The following morning before 9 am, females were removed from the males’ cages, and the ovaries with oviducts and upper uterine horns were dissected in one piece. The tissue was placed in a microtube with 1 ml of M2 medium and immediately transferred for processing. Under a stereomicroscope, zygotes surrounded by cumulus cells were isolated from the ampulla in a petri dish with M2 medium at 37°C. To remove the cumulus cells, zygotes were briefly put in hyaluronidase solution (HYASE-10X, Vitrolife) diluted 1:10 in M2 medium. The zygotes were then washed in three consecutive wells of 0.5 ml M2 medium covered with OVOIL-100 paraffin oil (Vitrolife). Normally fertilized zygotes, identified by the presence of two pronuclei, were selected and transferred to a pre-equilibrated micro-droplet culture dish (Vitrolife) containing 25 μl of G-1 PLUS culture medium (Vitrolife) per well, covered with paraffin oil, with approximately 10 zygotes per droplet. The zygotes were incubated at 37 °C with 6% CO₂ until developmental day 3.5, when they were further assessed. 8- cell embryos (n = 77), morulas (n = 11) and blastocysts (n = 14) had their zona pellucida removed using Tyrode’s acid treatment. The embryos were then washed in fresh M2 medium and cross- linked as described below (picoChIP-seq > *Oocyte and embryo cross-linking*).

### picoChIP-seq

The picoChIP-seq protocol has been described in detail elsewhere^24^ (manuscript in revision, available to reviewers upon request). A brief description of the different steps is provided:

#### Oocyte and embryo cross-linking

Oocytes were cross-linked in a solution of 50 µl M2 media, 50 µl PBS (Life Technologies, 14190-094), with a final concentration of 1% w/v formaldehyde (Sigma-Aldrich, F8775). After 8 min at room temperature, 14.3 µl 1 M glycine (Merck, 67419) was added to inactivate the fixation. Following 5 min at room temperature, ice cold PBS was used to wash the oocytes three times in a 0.6 ml tube (MaxymumRecovery, Axygen), resulting in a final volume of 10 µl. The fixative was added either in droplets under the microscope, or directly in the 0.6 ml tube followed by the washes and centrifugations at 700 x g for 10 min at 4°C with careful supernatant removal. Accordingly, the liquids were mixed by droplet pipetting or by gentle tube vortexing during cross-linking and glycine inactivation. Liquid nitrogen was used to snap-freeze the samples prior to storage at -80°C until further processing.

#### ES cell cross-linking

The mouse E14 embryonic stem cells (ESC) were cultured as described in the mouse ENCODE project instructions, starting from a stock at passage P2, equivalent to the cells used for the landmark project (https://www.encodeproject.org/biosamples/ENCBS171HGC/). Prior to collection, cells were passaged once on gelatinized, feeder-free plates. Approximately 20 million cells in a pellet were cross-linked at room temperature for 8 min, by resuspension in 20 ml of 1 % w/v formaldehyde (Sigma-Aldrich, F8775) solution in PBS, supplemented with 20 mM of sodium butyrate (Sigma-Aldrich, 19-137).The fixative was inactivated using 2.4 ml of an aqueous 1.25 M glycine (Sigma, G8790) solution, and the pellet was washed twice with ice cold PBS supplemented with 20 mM of sodium butyrate (650 x g, 10 min, 4°C). Following that, the cells were aliquoted into 0.6 ml tubes (MaxymumRecovery, Axygen) at various cell numbers. Finally, the cells were centrifuged, the supernatant removed, leaving 10 µl, that was snap-frozen in liquid nitrogen and stored at - 80°C.

#### Preparation of recombinant octamers

Plasmids for bacterial expression of recombinant human histones H2A, H2B, and H4 were generously provided by Robert Schneider (Helmholtz Zentrum, Munich, Germany) and H3.1 by Gunnar Schotta (LMU, Munich, Germany), and Histone octamers were prepared as reported previously^52^. The histones were expressed in *E. coli* BL21(DE3)pLysS, purified from inclusion bodies, and the pre-cleared extract was passed over a 5 ml HiTrap SP column. The four histones were reconstituted into octamers and separated using a HiLoad 16/60 Superdex 200 column in buffer (2 M NaCl, 10 mM Tris pH7.5, 1 mM EDTA pH 8.0, 5 mM b-mercapto-ethanol, and 0.2 mM PMSF). Equimolar fractions of the histones were pooled, concentrated, mixed with 50% glycerol (v/v), and stored at -20 °C.

#### Preparation of octamers and antibody-bead complexes

10 µg of recombinant octamers (2 µg/µl) were transferred to a 1.5 ml tube. These were cross-linked by addition of 5 µl of octamer buffer (2 M NaCl, 10 mM Tris-HCl pH 7.5, 1 mM EDTA pH 8.0) and 0.27 µl of octamer cross-linking buffer (3.7 % w/v formaldehyde in dH_2_O), resuspended by pipetting. Following a 20 min incubation at room temperature, the fixative was quenched for 5 min at room temperature using 0.72 µl of 200 mM glycine in PBS solution (Sigma-Aldrich, 67419-1ML-F), and stored at 4 °C. The octamers were used within three months (concentration was 0.9 µg/µl). Each picoChIP reaction was supplemented with 2.44 µg of the cross-linked octamers. Protein A Dynabeads (Invitrogen, 10002D) were incubated with antibodies a day prior to chromatin sonication. The beads were thoroughly vortexed and the volume specified in Table 2 was used for each picoChIP reaction and washed using RIPA buffer (10 mM Tris-HCl pH 8.0, 175 mM NaCl, 1 mM EDTA, 0.625 mM EGTA, 1.25% v/v Triton X-100, 0.125% w/v sodium deoxycholate, 1 x Halt-protease inhibitor cocktail (ThermoFisher, 78440), 1 mM PMSF, 20 mM sodium butyrate) two times. To improve handling, an excess of beads was washed using a PCR-strip magnetic rack (Diagenode), a 0.6 ml tube magnetic rack (manufactured in-house), and a hand-held strong neodymium magnet for complete separation when needed. The master mix of washed beads was prepared in a suitable tube (1.5 ml / 0.6 ml / 0.2 mL, Axygen), taking into consideration the intended number of ChIP reactions and the volumes given in Table 2, followed by dilution in 20-100 µl RIPA per reaction. The corresponding volume of antibody (Table 2), followed by incubation on a ‘head-over-tail’ rotator (10 rpm) at 4 °C overnight (ON).

**Table 2.** Antibody information for H2A.Z picoChIP.

| Table 2. Antibody information for H2A.Z picoChIP |  |  |  |  |
| --- | --- | --- | --- | --- |
| Epitope | Antibody cat. #no. | Antibody lot. | Bead stock (μl) | Antibody (μl) |
| H2A.Z | 39114, Active Motif | 06217001 | 2 | 2 |

#### Chromatin preparation

On the following day, transfer of the cross-linked samples was done using dry ice and thawing in a cold block for a few minutes, after which 120 µl lysis buffer (50 mM Tris–HCl pH 8.0, 10 mM EDTA pH 8.0, 0.8% w/v SDS, 1 x protease inhibitor cocktail, 1 mM PMSF, and 20 mM sodium butyrate) were added. Following 5-15 minutes incubation of the samples in a cold block of approximately 6-8°C, 30 µl of PBS supplemented with 20 mM sodium butyrate were added. Then, the samples were sonicated as per Table 3, using a Hielscher UP100H sonicator equipped with a 2 mm probe, configured at 27% amplitude and 0.5 s pulse intervals.

**Table 3.**
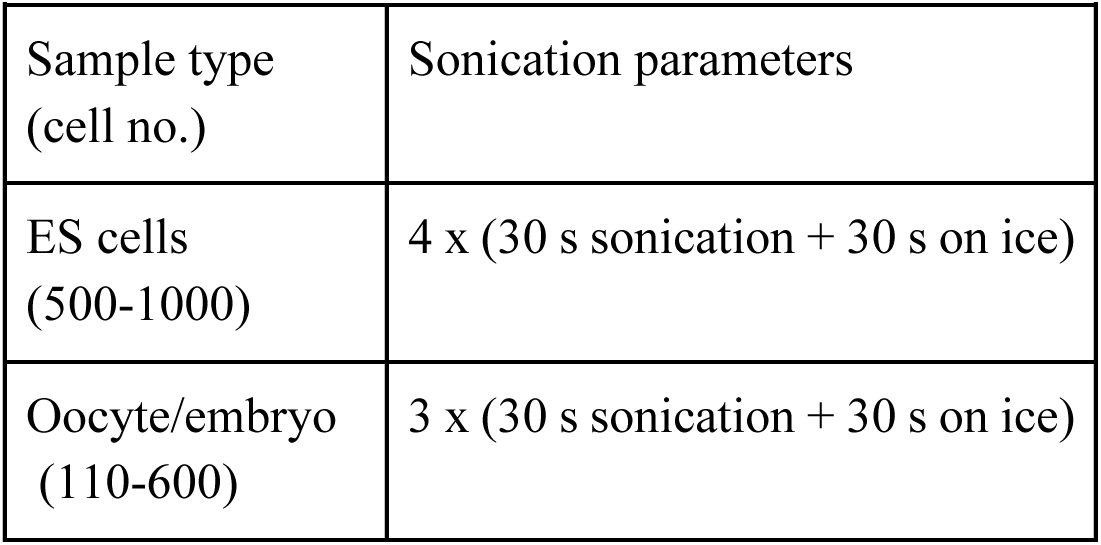
Sonication parameters for picoChIP.

Directly after sonication, the tubes containing 160 µl of sonicated chromatin were diluted with the appropriate volume RIPA dilution buffer (10 mM Tris-HCl pH 8.0, 175 mM NaCl, 1 mM EDTA, 0.625 mM EGTA, 1.25% v/v Triton X-100, 0.125% w/v sodium deoxycholate, 1 x protease inhibitor cocktail, 1 mM PMSF, and 20 mM sodium butyrate) and processed further as per Table 4.

**Table 4.**
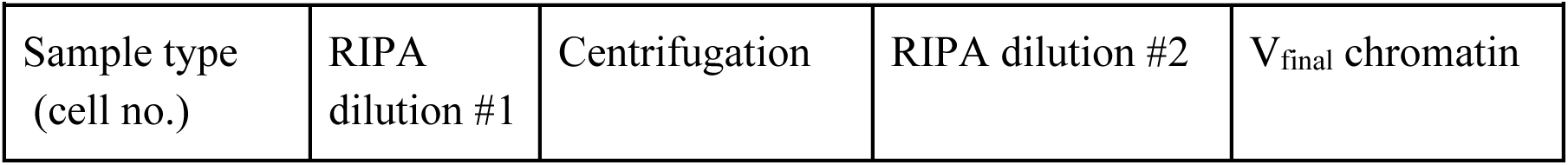
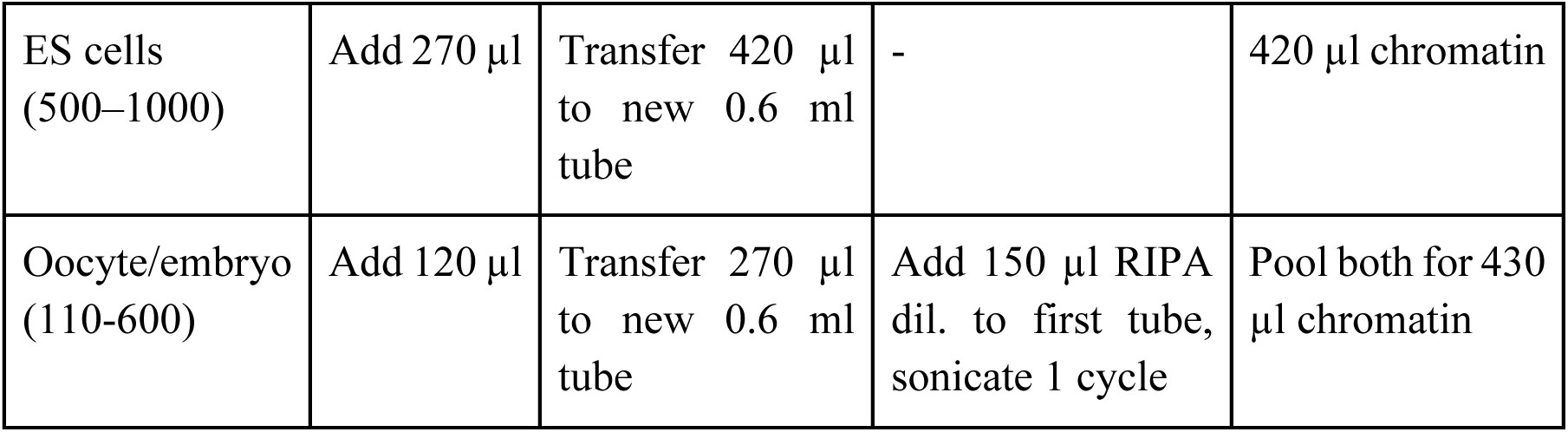
Sonicated chromatin preparation for H2A.Z picoChIP.

Centrifugations were performed at 16,000x *g* for 10 min, at 4 °C. At this point, the 5 % input samples were acquired from the 430 µl chromatin by storing 21.5 µl at 4 °C. Chromatin preparations intended for multiple ChIP reactions were divided in fractions of equal volume, containing the desired cell number chromatin equivalents, and the volume to a final of 430 µl reached using volume adjustment buffer (40 parts PBS with 20 mM sodium butyrate, 120 parts lysis buffer, 270 parts RIPA dilution buffer).

#### Chromatin immunoprecipitation reaction setup

The antibody-bead complexes were washed with RIPA buffer and robust vortexing three times, and the coated beads were resuspended in a total of 102 µl RIPA dilution per picoChIP reaction, of which 100 µl were added to each 430 µl sonicated chromatin aliquot. Each picoChIP reaction was supplemented with 1.25 µl IgG (Abcam, ab46540) and 2.44 µg of the cross-linked octamers, and incubated on a ‘head-over-tail’ rotator at 10 rpm for 30-40 hours at 4 °C.

#### Washes and elution

Following the extensive incubation, the tubes were vortexed thoroughly, any droplets collected by a short centrifugation, the beads pelleted with a hand-held magnet, the supernatant removed, and the beads were transferred to 0.2 ml strip tubes using 150 µl of the first wash buffer. The washing steps for H2A.Z picoChIP-seq are given in Table 5.

**Table 5.** Sample washing steps for H2A.Z picoChIP.

| Table 5. Sample washing steps for H2A.Z picoChIP |  |  |  |  |
| --- | --- | --- | --- | --- |
|  | Wash 1 | Wash 2 | Wash 3 | Wash 4 |
| Volume (μl) | 150 | 100 | 100 | 100 |
| Buffer | RIPA-standard | RIPA-high | RIPA-high | RIPA-standard |

The employed buffers provided wash steps of different stringency due to their compositions: <u>RIPA-standard</u> (10 mM Tris-HCl pH 8.0, 140 mM NaCl, 1 mM EDTA, 0.5 mM EGTA, 1% v/v Triton X-100, 0.1% w/v SDS, 0.1% sodium deoxycholate, 1 x protease inhibitor cocktail, 1 mM PMSF, 20 mM sodium butyrate) and <u>RIPA-high</u> (10 mM Tris-HCl pH 8.0, 300 mM NaCl, 1 mM EDTA, 0.5 mM EGTA, 1% v/v Triton X-100, 0.22% w/v SDS, 0.1% v/v sodium deoxycholate, 1 x protease inhibitor cocktail, 1 mM PMSF, 20 mM sodium butyrate). A magnetic rack was used to perform the washes. For each washing step, stringency was increased mechanically via vortexing for 3 x 5 s on the highest setting, repeated twice, with 10 s incubation on ice in between. Following the last washing step, TE buffer (10 mM Tris–HCl pH 8.0, 1 mM EDTA pH 8.0) was used to transfer each picoChIP sample to a new 0.2 ml tube, and the beads were eluted using 150 µl of elution buffer (20 mM Tris–HCl pH 7.5, 50 mM NaCl, 5 mM EDTA pH 8.0, 1% w/v SDS and 0.2 mg/ml RNase A). At this point, the 5% input chromatin tubes were mixed with 128.5 µl elution buffer, and were treated as the other samples moving forward. RNase incubation was performed for 1 h at 37 °C with vigorous agitation (1250 rpm), followed by addition of 1 µl of Proteinase K (20 mg/mL, NEB, P8107S) and incubation for 4 hours at 68 °C with vigorous agitation.

#### DNA purification, library preparation and quantification

The picoChIP and input samples were transferred to 1.5 ml tubes. SDS/RNase-free elution buffer (20 mM Tris–HCl pH 7.5, 50 mM NaCl, 5 mM EDTA pH 8.0) was added to 415 µl final volume, and the aqueous phase was sequentially extracted using equal volumes of phenol:chloroform:isoamylalcohol (Invitrogen, 15593-031) and chloroform:isoamylalcohol (Sigma-Aldrich, C0549). To the 400 µl of the last aqueous phase 1 ml ice-cold ethanol, as well as 11 µl linear acrylamide (Thermo Fisher Scientific, AM9520) and 44 µl 1 M sodium acetate (Invitrogen, AM9740) were added to precipitate the DNA at -80°C at least overnight. For reconstitution, the pellets were centrifuged (swing-out buckets, 16,000x *g*, 4 °C), and the DNA recovered using 15 µl of Qiagen EB (10 mM Tris, Qiagen, 19086) overnight at 4 °C. Illumina sequencing libraries were prepared using the Qiaseq ultralow input kit (Qiagen, 180492 + 180310), optimized to minimize material losses. These included prolonged incubations during the recoveries from AMPure XP beads (Beckman coulter, A63881), with >15 min bead/sample incubation, and >15 min DNA elution, as well as an increased volume of beads (1.2 volume equivalents) during the second bead extraction. Double-stranded DNA concentration was determined using the Qubit HS DNA kit (ThermoFisher Scientific, Q32851), while DNA fragment size analyses were performed using the TapeStation D1000 HS kit (Agilent, 5067-5585). Equal moles of each library were pooled and sequential AMPure XP extractions (1.2 volume equivalents) reduced adapter dimers from the pool prior to sequencing on a NovaSeq 6000 SP, paired end 50 bp, at the Norwegian Sequencing Center (Ullevål, Oslo).

### Data analysis

#### Data sources

All new ChIP-seq data are deposited at NCBI’s Gene Expression Omnibus^53^ under the accession number GSE293415. Datasets of H3K4me3, H3K27me3, and DNA methylation in mouse oocyte and early embryo development was sourced from^31, 33, 54, 55^ and downloaded from GEO^53^ series GSE72784; GSE56879; GSE73952 and GSM2041068. RNA seq data^27, 32^ were downloaded from ArrayExpress^56^ (Accession Number P-MTAB-41287), and GEO^53^ GSM1845293. Aggregated single-cell Repli-seq data of the maternal and paternal genomes in 2- cell mouse embryos were processed in^34^ and obtained from Supplementary Data 7 in the paper. The mean Replication Status values from all single-cells were used for analysis and visualization here. Bed-files with allele-specific coordinates for LADs in maternal and paternal 2-cell embryos^21^ were downloaded from the NCBI GEO database^53^ series GSE112551. Refseq gene annotations for mm10 were downloaded from the UCSC table browser^57, 58^ and coordinates for unique TSSs were defined based on this (n=32,166). CGIs as defined in the mouse mm10 genome were also downloaded from the UCSC table browser^58^. CpG and GC densities from the mm10 reference genome sequences, were downloaded from the UCSC genome browser^58, 59^ at https://hgdownload.soe.ucsc.edu/goldenPath/. A mouse enhancer set titled ENC+EPD enhancers (n=37,473) was sourced from the UCSC table browser^36, 58, 60^. A dataset of different embryonic gene categories in the mm10 genome was sourced from DBTMEE v2 transcriptome categories^61^. A dataset of LTR-initiated transcription units (LITs) (n=3384) was obtained from^39^. Repeat sequences from the mouse genome was downloaded from the UCSC table browser^58^ (Repeatmasker 21-10-2024^37^).

#### ChIP-seq and RNA-seq data processing

ChIP-seq and RNA-seq data was processed using an in-house Nextflow pipeline (https://github.com/lerdruplab/ew-qctrimalign) written according to nf-core guidelines^62^ utilizing reproducible software environments from Bioconda^63^ and Biocontainers^64^. Briefly, the pipeline was executed using Nextflow v23.4.1^65^. Single-end fastq files were trimmed using Trim Galore v0.6.7 (https://github.com/FelixKrueger/TrimGalore) with the parameter -novaseq 20. Trimmed reads were then mapped to mm10 using bowtie v1.3.0^66^ with the parameter -m 1. Obtained bam files were sorted using samtools v1.2^67^ and converted to bed format using bedtools v2.31.1^68^. Bed files were then used for downstream analysis in EaSeq^69, 70^.

#### General data visualization

Unless stated, values are FPKM normalized. Simple plots, including doughnut plots, scatter plots, line plots and tiles were generated in Microsoft Excel or using R^71^. Bubble plots were made using R^71^ by calculating the log_2_fold difference between observed and expected number for each analyzed feature, Chi Square tested to assess significance for each combination, and adjusted for multiple statistical tests using the Benjamini-Hochberg procedure to control the false discovery rate (FDR) (Github repository: https://github.com/lerdruplab/H2A.Z). The following tools in Easeq^69, 70^ were used for data visualization and adjusted as shown in the figures: Genome browser tracks were generated using the “FillTrack” tool, Metagene plots were generated using the “Average” tool followed by the “Overlay” tool, Colored 2D-histograms were created with the “ZScatter” tool, and heatmaps and simple heatmaps were generated using the “HeatMap” and “Parmap tools, respectively. Overlap between features was calculated using the “Coloc” tool with the setting “Distances measured from border to border of the regions”. For visualization of features throughout the mouse genome, the “Modify” tool was used to subdivide (“Homogenize”) a regionset listing sizes for each chromosome in the mm10 genome into 1 kbp bins (n=2,633,787) and 10 kbp bins (n=263,389), respectively.

#### Clustering

Regions were clustered in Easeq^70^ using the “Cluster” tool set to perform to k-means clustering using a k-value of 10. Initial k-means were assigned using the k^++^ approach^72^. Clustering start point, end point, and offsets for each analysis is stated at the respective analysis and figure.

#### H2A.Z peak calling and aggregation

Peaks were called from pooled replicate samples of developmental stages individually against corresponding input samples and then merged into one peak set (n=111,448, Table S1) in Easeq with default parameter settings as described in^70^ using “Regionsets/Modify/Merge” in a consecutive manner. We then quantified all samples at the center of the peak +/-1kbp using the “Quantify” tool in EaSeq. For clustering, quantified values based on H2A.Z from all developmental stages in the aggregate peak set were quantile normalized to ensure that H2A.Z signal from all samples contributed equally prior to clustering using the “Normaliz.” and “ClusterP” tools in EaSeq, respectively. A principal component analysis of the peaks was carried out in R using the bioconductors “pcaMethods” package with default parameters^71, 73^, to visualize the variance between samples (Github repository: https://github.com/lerdruplab/H2A.Z). To visualize enrichment/depletion of CpG density, H3K4me3 and H3K27me3 histone marks^31, 33, 54^ in peaks compared to genome-wide signal, Z-scores were calculated in the complete clustered peak set as well as the non-TSS/CGI peak set using the “Quantify” tool in Easeq. When replicates were available, average Z-scores were calculated prior to visualization.

#### H2A.Z peak annotation

Peak annotation was performed using ChIPseeker v1.38.0^74^ in R v4.3.3. Gene ontology analysis of H2A.Z peak clusters were performed in R v4.3.3 using gprofiler2^75^ and visualized as a heatmap using ComplexHeatmap^76^. We used the full-stack ChromHMM segmentation data for the mm10 mouse assembly (<u>github.com/ernstlab/mouse_fullStack_annotations</u>^26^ to examine overlap of the annotation to the complete H2A.Z peak set and the non-TSS/CGI peak set. Using shuffle bed from BEDtools^68^, five randomized control regions of equal number and length to the peaks were created, and the overlaps counted with annotate bed also from BEDtools. We depicted the average relative enrichment with the standard deviation as error bars. Additionally, gene assignments for the peaks were done by ChIP-Enrich (chip-enrich.med.umich.edu^77^) and the statistically significant GO:BP genesets (FDR ≤ 0.05) were processed by Revigo (revigo.irb.hr^78^). Cytoscape was used to visualize the resulting networks (cytoscape.org)^79^.

#### H2A.Z at TSSs and CGIs

For subgrouping, H2A.Z signal at TSSs at the individual stages was quantified and FPKM normalized at the center of the peak +/-1kbp using the “Quantify” tool in EaSeq, and clustered using the “ClusterP” tool in EaSeq. Total RNA expression data obtained from^27^, were quantified and FPKM normalized at TSSs (0 +/-500 bp) then quantile normalized using “Normaliz.” tool in EaSeq. H2A.Z signal and quantile normalized total RNA signal was then visualized as heatmaps based on the clustering. To find enriched or depleted gene categories for each cluster, we calculated the number of genes from different embryonic transcriptome categories (DBTMEE v2 transcriptome categories^61^) in each cluster and visualized in Bubble plots as described above. To explore the relationship between H2A.Z, CGIs and other genomic features at TSSs, CGIs were colocalized to TSSs using the “Coloc.” tool in EaSeq. Calculated distances where sorted based on proximity to the nearest CGI using the “Sort” tool. Correlations of H2A.Z, H3K4me3^31^, and CpG densities both at TSSs (n=32,166) and in the whole genome divided into 1 kbp bins (n=2,633,787) were calculated with Spearman’s rank correlation coefficients in R^71^.

#### H2A.Z at repeats

To analyze the H2A.Z signal at repeats, we colocalized H2A.Z peaks with repeat coordinates, and subselected repeats with a count of more than 200 occurrences in the aggregated peak set to ensure solid statistical analyses. To find enriched or depleted repeat categories for each cluster, repeat coverage as well as abundance was calculated for each peak cluster in the non-TSS/CGI peak set and used for visualization. Selected repeat types were chosen based on high abundance in clusters 1 and 2, and the P12 stage was used to visualize H2A.Z signal at these. Repeat types were clustered at the start of the repeat +/-3kbp and sorted based on the mean H2A.Z signal in each cluster. Given the high H2A.Z signal at MTA RTs, LITs from^39^ were sorted based on overlap to CGI and H2A.Z peaks, and used for visualization of local H2A.Z, H3K4me3 and H3K27me3 signals^31, 33, 54^.

#### Motif enrichment analysis and integration with CpG / G/C density data

Identification of enriched DNA motifs were performed on aggregate H2A.Z peaks, which were not overlapping with TSSs or CGIs, from oocyte-specific clusters (1, 2 and 3) and embryo-specific clusters (8,9 and 10). Sets of random control regions with matching distances to the nearest TSS were generated using the “Controls” tool in EaSeq, and sequences from the peaks and control regions were extracted using the “Get Sequences” tool in EaSeq (in the beta testing menu), and used for de novo motif enrichment analysis using the program MEME-ChIP^80^. Only hits with E-value ≤ e^-10^ were considered.

## Acknowledgements

We thank the members of the Lerdrup group, Dahl group, CRESCO, and the Center for Chromosome Stability for support and critical suggestions throughout this work. In addition, we thank the Norwegian Sequencing center, The Norwegian Transgene center, and the staff at the animal facilities (Department of Comparative Medicine, Oslo University Hospital and Department of Comparative Medicine, University of Oslo). This work was supported by the Danish National Research Foundation DNRF115 (M.L), the Novo Nordisk Foundation, NNF22OC0080710 (M.L.), and Norwegian Centres of Excellence scheme grant 332713 - CRESCO (M.L. and J.A.D.).

## Author contributions

M.F., M.L. and J.A.D. conceived the study. M.F. designed the experiments with input from J.A.D. M.F. carried out all picoChIP-seq experiments. Generation of recombinant octameres used in picoChIP-seq was carried out by R.E. Mouse work and ovary collection was performed by T.S., I.J. and S.K. under supervision by G.G., K.T.D. P.F. and J.A.D. Mouse oocyte and embryo collections were planned and performed by M.F., T.S., A.M., R.S., M.B., M.V-R. and J.A.D. E.I., M.S., M.C and M.Z. performed the data analysis with input, assistance and supervision from M.L. M.F., E.I., M.Z., M.C, M.L. and J.A.D. discussed the data and provided critical input. M.Z., E.I, and M.L. prepared the manuscript with assistance from M.F., G.G., P.F., T.S., M.S, and J.A.D. All authors read and commented on the manuscript.

## Supplementary information

The online version contains supplementary material available at

## Data availability

All new ChIP-seq data are deposited at NCBI’s Gene Expression Omnibus^53^ under the accession number GSE293415.

## Ethics declarations

All mouse experiments were approved and registered by the Norwegian Food Safety Authority (NFSA approved application/FOTS IDs: 7216, 10898, and 24911, and 8743) and conducted in accordance with Norwegian regulation FOR-2015-06-18-761, which closely aligns with EU directive 2010/63/EU on stringent ethical and welfare standards to protect animals used for scientific purposes.

## Supplementary information figure legends

**Fig. S1.**
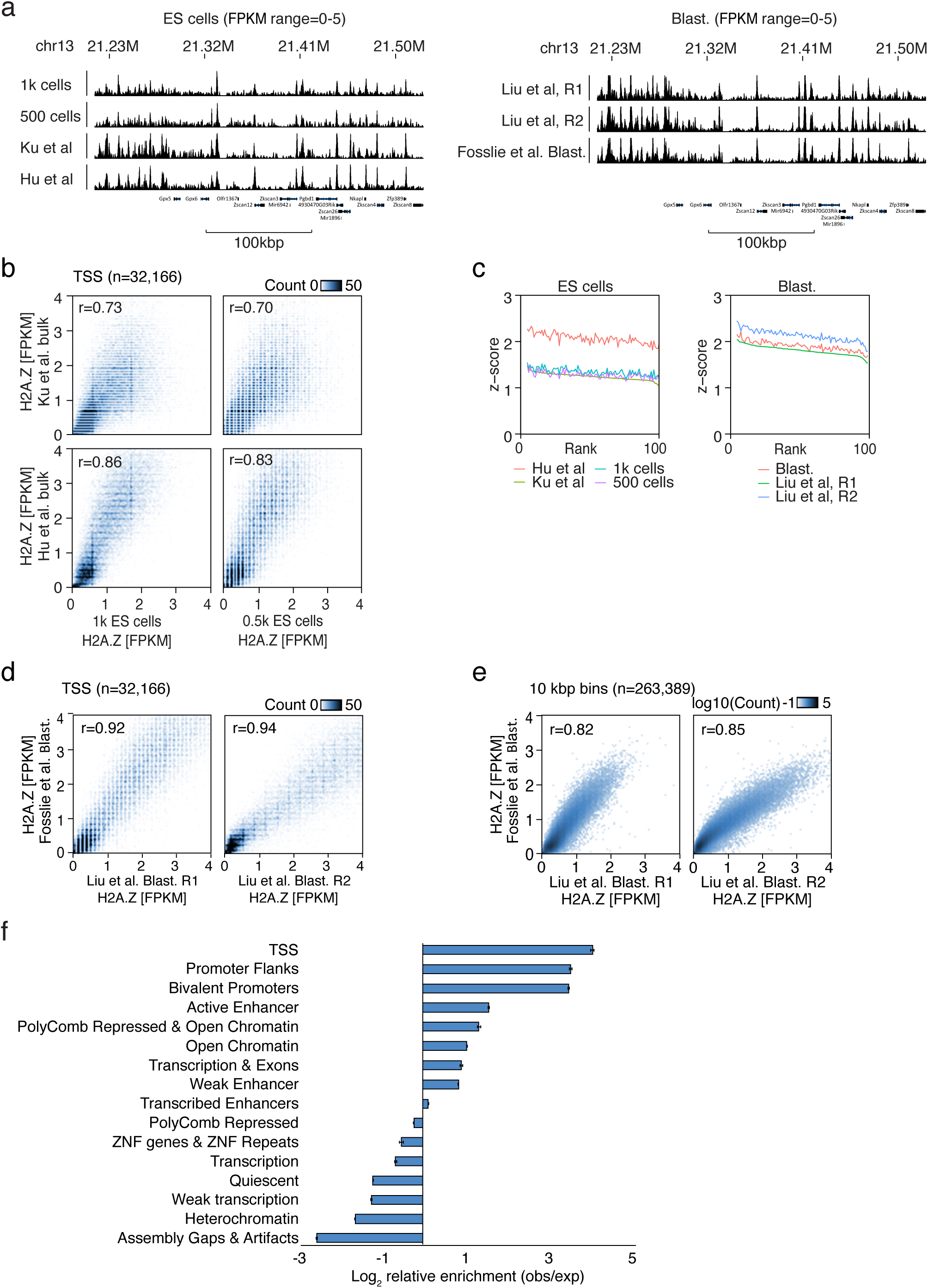
H2A.Z samples from this study showed good correlations compared to previously published data. **a**, Genome tracks of H2A.Z ChIP-seq signal from this study compared to previously published H2A.Z ChIP-seq signal from ES cells^13, 25^ and blastocysts (blast.)^14^. **b**, 2D-histograms showing the relationship between H2A.Z enrichment in ES cells from this study to ES cells from^13, 25^ at TSSs (n=32,166) +/-1kbp. r: Spearman’s rank correlation coefficients. **c**, Z-scores of H2A.Z signal at TSSs (+/- 1kbp) in ES cell^13, 25^ and blastocyst samples^14^. Z-scores are ranked based on^14, 25^ R1 samples in the respective plots. **d**, **e**, 2D-histograms showing the relationship between H2A.Z enrichment in blastocysts (blast.) from our study (Y-axis) and previously published samples (X- axis)^14^ at unique TSS (**d**) (n=32,166) +/-1kbp, and throughout the genome divided into 10 kbp bins (**e**) (n=263,389). r: Spearman’s rank correlation coefficients. **f**, Analysis of H2A.Z peaks in functional chromatin regions conserved across mouse tissues^26^. The average relative enrichment is shown with the standard deviation as error bars.

**Fig. S2.**
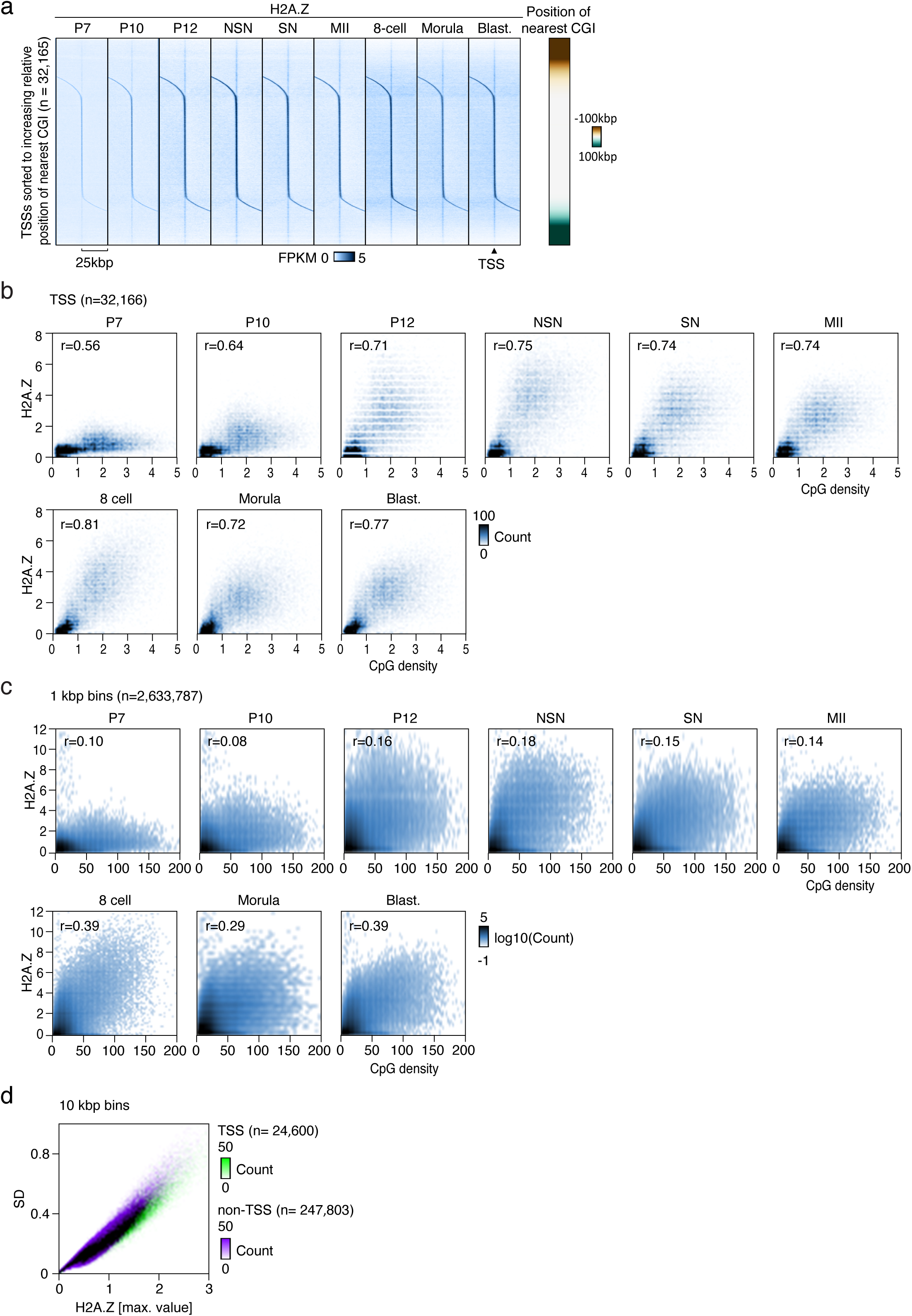
H2A.Z is correlated to CpG density at TSSs but not in the overall genome. **a**, Heatmaps of H2A.Z signal in different developmental stages at unique TSSs +/-25kbp (n=32,166) sorted according to increasing relative position of the nearest CGI. **b**, **c**, 2D-histograms showing the relationship between H2A.Z and CpG density in different developmental stages at (**b**) unique TSSs (n=32,166) +/-1kbp and (**c**) throughout the whole genome (1kbp bins, n=2,633,787). r: Spearman’s rank correlation coefficients. **d**, Overlaid 2D-histograms showing genome-wide relationships between maximum H2A.Z levels (X-axis) as well as standard deviations (Y-axis) of all stages analyzed in 10 kbp bins (n=263,389). Purple and green coloring represents bins that do and do not overlap with a TSS, respectively. Black coloring represents the overlay between these two populations.

**Fig. S3.**
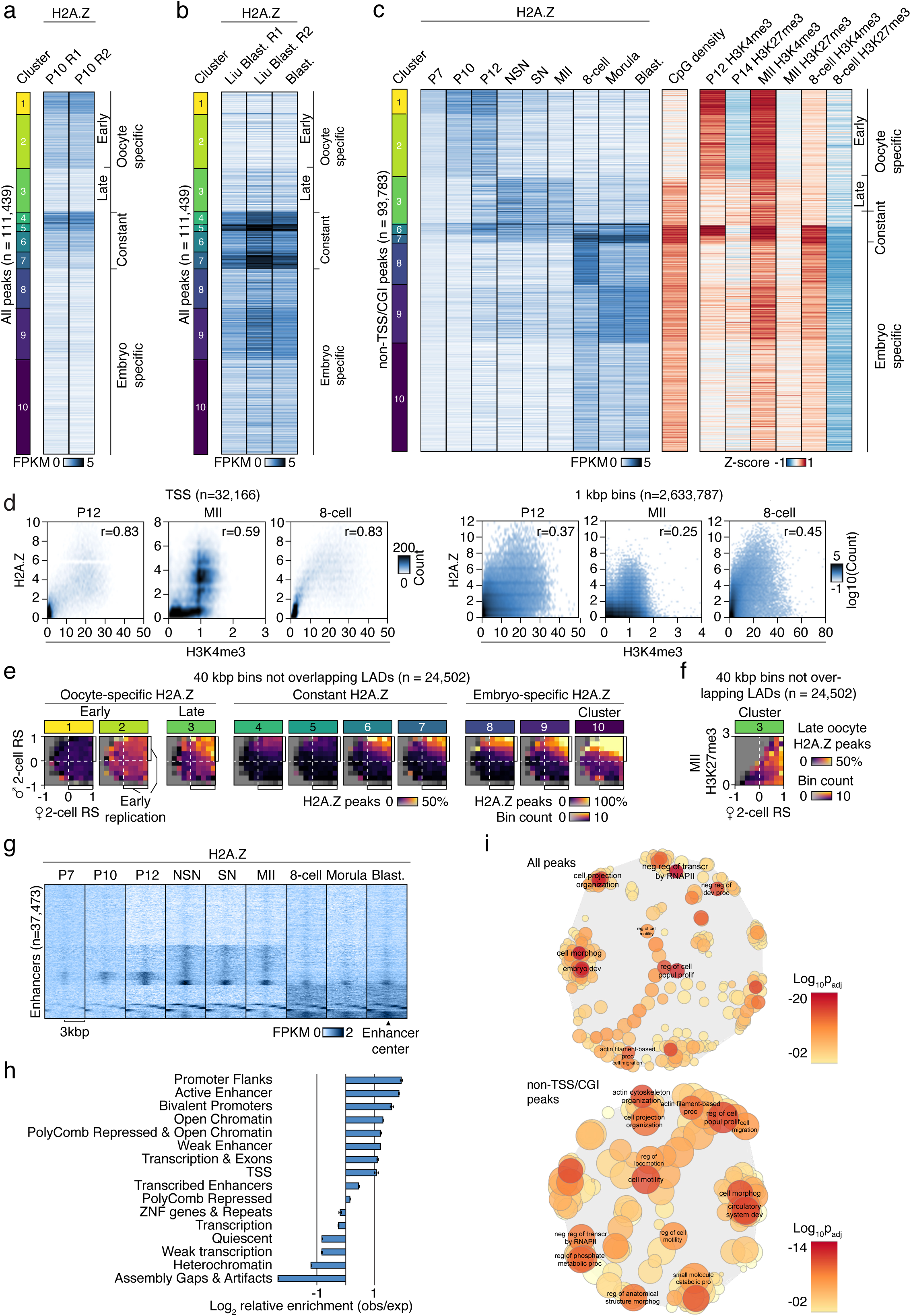
H2A.Z peaks are correlated to H3K4me3 at TSSs but not in the overall genome. **a**, Heatmaps of H2A.Z enrichment at aggregated and clustered H2A.Z peaks at two P10 replicates from this study. **b**, Heatmaps of H2A.Z enrichment at aggregated and clustered H2A.Z peaks in blastocyst (blast.) replicates from^14^ compared to blastocyst from this study. **c**, Heatmaps of H2A.Z enrichment at aggregated and clustered H2A.Z peaks during different developmental stages as well as Z-score of CpG density and histone marks^31, 33, 54^ in clustered peaks (non-TSS/CGI, n=93,783) compared to genome-wide signal. **d**, 2D-histograms showing the relationship between H2A.Z and H3K4me3^31^ signal at selected stages at either TSSs (n=32,166) +/-1kbp or whole genome (1kbp bins, n=2,633,787). r: Spearman’s rank correlation coefficients. **e**, 2D-histograms showing the genome-wide occurrence of each cluster of H2A.Z peaks (color) in relation to the mean Replication Status of the maternal (X-axis) and paternal (Y-axis) genomes^34^ in individual 2-cell embryos. # and * indicates noteworthy sex-specific differences, where low and high peak- occurrences, respectively, largely follows the maternal Replication Status, but not the paternal. Data were analyzed in 40 kbp bins, and the subset of bins not overlapping with LADs are shown (n=24,502). For the overlapping subset see Fig 3d. **e**, 2D-histogram showing the genome-wide occurrence of cluster 3 H2A.Z peaks (color) in relation to the mean maternal Replication Status (X-axis) in individual 2-cell embryos and H3K27me3 levels in MII oocytes (Y-axis). Data were analyzed in 40 kbp bins, and the subset of bins not overlapping with LADs are shown (n=24,502). For the overlapping subset see Fig 3d. **g**, H2A.Z signal at enhancers (n=37,473) in different developmental stages, clustered based on the stage-specific distribution of the H2A.Z signal at and around the enhancers. **h**, Analysis of H2A.Z non-TSS/CGI peaks in functional chromatin regions conserved across mouse tissues^26^. The average relative enrichment is shown with the standard deviation as error bars. **i**, Networks of enriched GO terms in all and non-TSS/CGI H2A.Z peak sets show significant enrichment in genes involved in transcriptional regulation, cell morphogenesis, and developmental processes. H2A.Z peaks were assigned to gene sets, the enriched processes (FDR ≤ 0.05) summarized by REVIGO^78^, and plotted in Cytoscape. Exemplary GO terms for each cluster are annotated. Semantic similarity reflected in the clustering, color and font size reflect the log_10_ *FDR* value. Circle radius reflects the log of number of genes in GO term ID. dev, development; morphog, morphogenesis; neg, negative; popul, population; proc, processes; prolif, proliferation; reg, regulation; transcr, transcription.

**Fig. S4.**
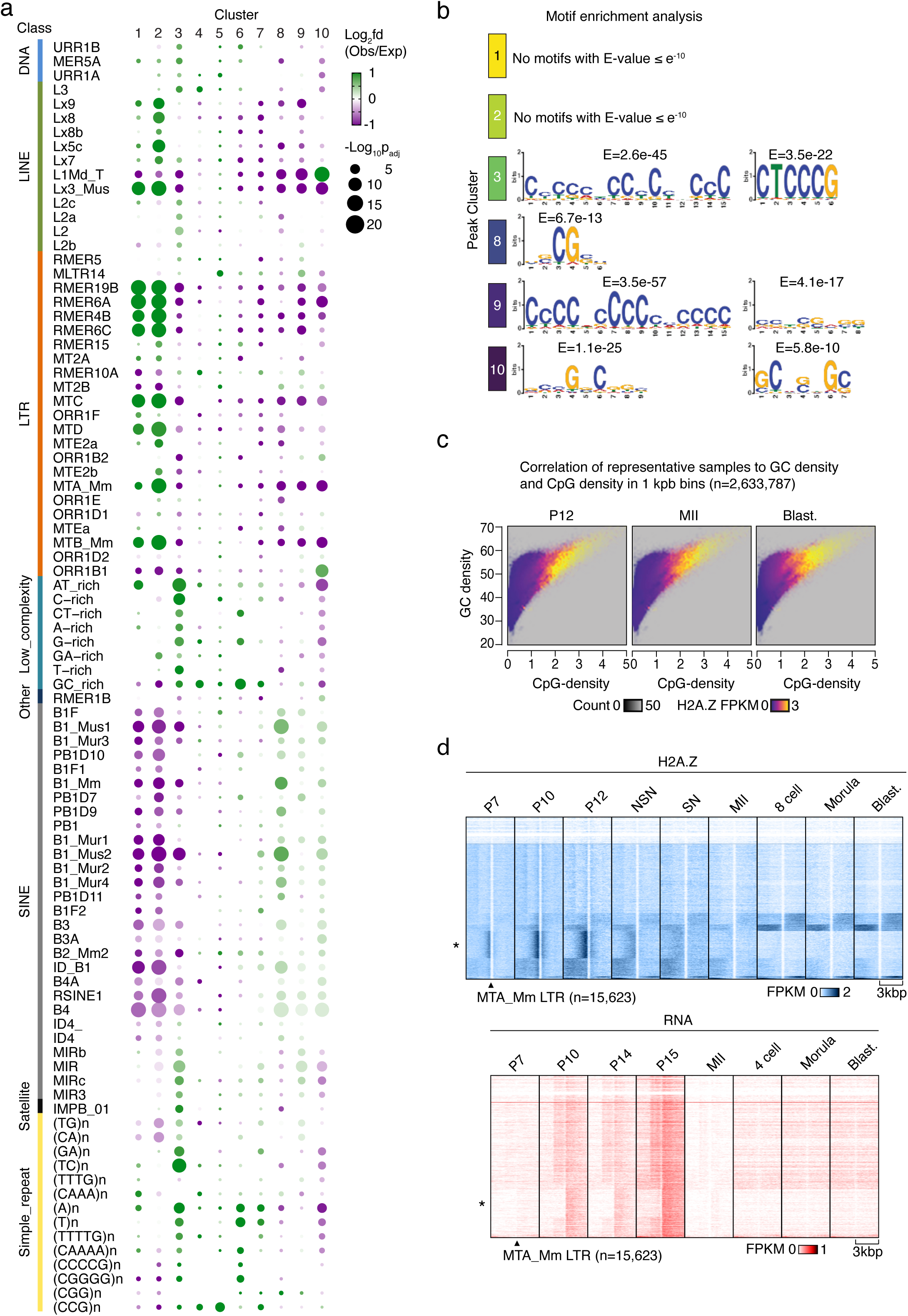
H2A.Z is correlated to CpG density and found at specific RTs. **a**, Bubble plots showing overlap of repeat subtypes in clusters of non-TSS/CGI H2A.Z peaks compared to an average distribution across all clusters. p-values are calculated by Chi Square tests Benjamini-Hochberg adjusted for multiple testing. **b**, Graphical presentations of nucleotide composition and frequencies in the most enriched motifs at selected clusters of aggregate H2A.Z peaks found using MEME-ChIP^80^. Peaks were identified and clustered as Fig. 3, and peaks from oocyte specific clusters (1, 2 and 3) and embryo specific clusters (8,9 and 10) not overlapping with TSSs or CGIs were tested. Control regions were based on distance to the nearest TSS, and only hits with E-value ≤ e^-10^ are shown. **c**, 2D-histograms showing the relationship between GC density, CpG density and H2A.Z signal from selected developmental stages throughout the whole genome (1 kbp bins, n=2,633,787). **d,** Heatmaps of H2A.Z and RNA signal^27^ surrounding the start of MTA LTR RTs in different developmental stages clustered based on H2A.Z in all stages. *Highlights cluster with the strongest H2A.Z signal in P12 oocytes.

**Fig. S5.**
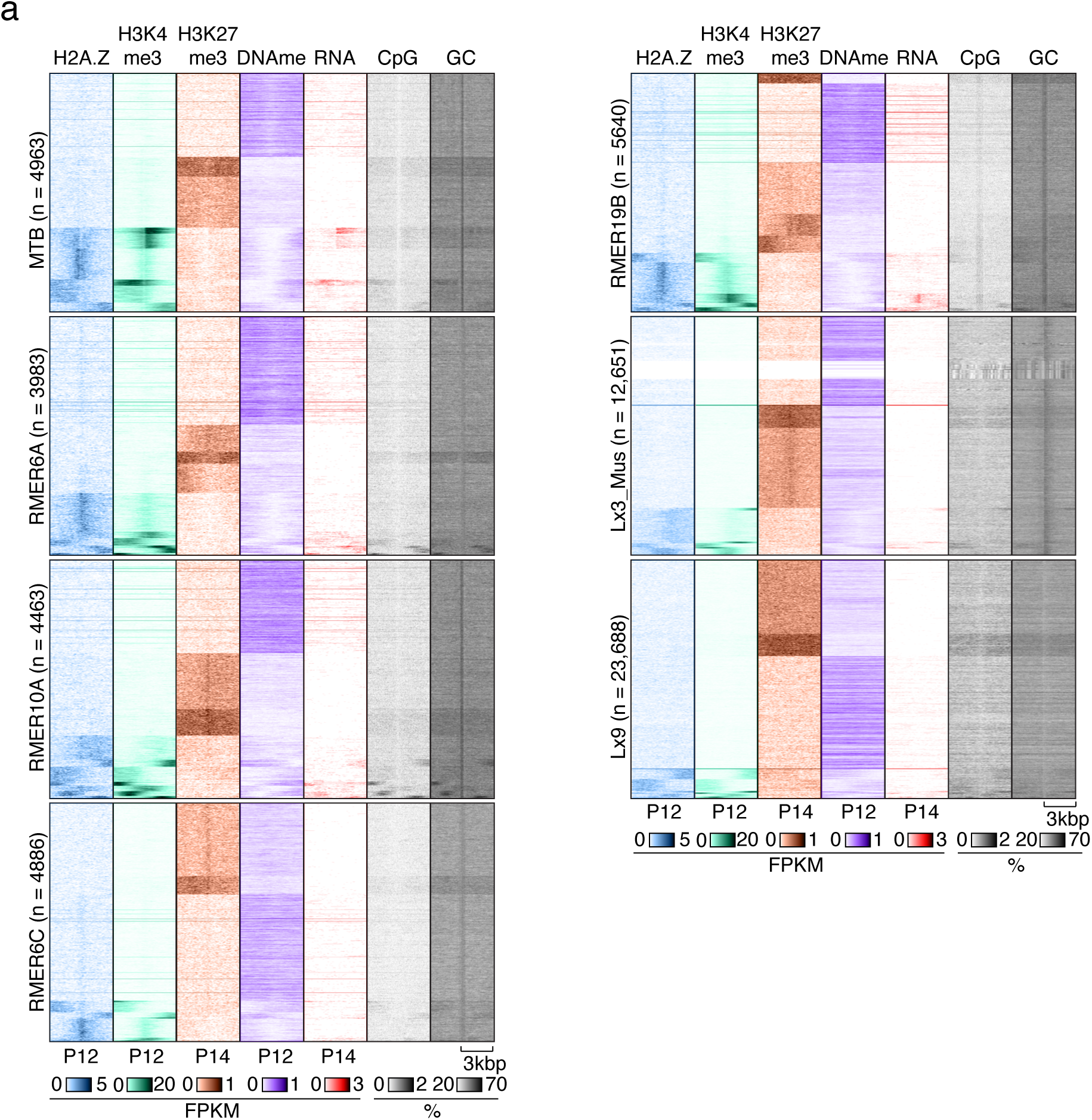
H2A.Z peaks overlap with specific repeats. Heatmaps showing indicated features^27, 55^ at selected RTs clustered based on the combined distribution of H2A.Z, H3K4me3^31^ and H3K27me3^54^ at the start of each RT +/-3kbp in growing oocytes (P12 or P14).

