## Supplementary Table 2 for "Major waves of H2A.Z incorporation during mouse oogenesis"

| go_term:bp | c1_-log10_padj | c2_-log10_padj | c3_-log10_padj | c4_-log10_padj | c5_-log10_padj | c6_-log10_padj | c7_-log10_padj | c8_-log10_padj | c9_-log10_padj | c10_-log10_padj |
|---|---|---|---|---|---|---|---|---|---|---|
| positive regulation of biosynthetic process | 0 | 0,318757668 | 8,251635988 | 17,52510183 | 22,96733842 | 34,05747214 | 37,5681275 | 11,83033562 | 23,90376844 | 14,43770229 |
| regulation of signal transduction | 0 | 0 | 20,94341703 | 8,585300875 | 42,95986405 | 22,93073816 | 46,22279384 | 8,026775661 | 18,70296637 | 14,49174653 |
| cellular process | 0 | 0 | 13,36156464 | 80,34429594 | 34,49794137 | 79,77812605 | 31,47579238 | 14,42877083 | 14,63473384 | 12,37277413 |
| metabolic process | 0 | 0 | 0 | 69,76677848 | 16,72326946 | 56,40953618 | 6,472302204 | 3,107980673 | 4,290768736 | 1,015861094 |
| cellular component organization | 0 | 0 | 15,06848396 | 58,21835208 | 32,45662905 | 50,93606157 | 24,58990916 | 9,443974431 | 9,352731874 | 9,263014787 |
| positive regulation of macromolecule biosynthetic process | 0 | 0 | 7,916789288 | 17,46483659 | 22,59011399 | 31,24329709 | 34,99090949 | 11,56258486 | 22,28144787 | 13,22624061 |
| positive regulation of macromolecule metabolic process | 0 | 0 | 11,54379457 | 29,35229925 | 33,87735218 | 44,1868629 | 41,22656155 | 12,29002615 | 26,58436922 | 17,90831383 |
| regulation of cell communication | 0 | 0 | 25,83715855 | 5,818355607 | 48,32459455 | 21,19801015 | 51,84413143 | 7,554391632 | 19,81304588 | 11,4589917 |
| DNA-templated transcription | 0 | 0 | 3,576124244 | 43,66091599 | 41,30631835 | 29,95411008 | 35,6415665 | 7,788746029 | 16,12052932 | 7,614375171 |
| regulation of DNA-templated transcription | 0 | 0 | 4,076962047 | 38,62250298 | 43,86174343 | 26,60928259 | 38,93934069 | 8,241166227 | 16,87756125 | 8,272782608 |
| regulation of transcription by RNA polymerase II | 0 | 0 | 5,041564992 | 29,07609587 | 37,2377878 | 26,20156707 | 45,59331229 | 10,36821729 | 19,87706266 | 12,04777191 |
| transcription by RNA polymerase II | 0 | 0,404606085 | 4,117716238 | 33,4499798 | 35,43160592 | 28,77835135 | 42,3450415 | 10,16350765 | 19,08819481 | 11,65328637 |
| protein modification process | 0 | 0 | 4,19341712 | 45,19273228 | 38,9801086 | 42,11997732 | 18,32569146 | 6,209189751 | 10,2758795 | 6,956119262 |
| intracellular signal transduction | 0 | 0,758332175 | 24,27461581 | 13,1910373 | 35,47897947 | 32,99630451 | 39,59245699 | 9,694381365 | 17,25470649 | 18,66900214 |
| regulation of signaling | 0 | 0 | 25,01204483 | 5,98048089 | 47,97012318 | 21,67977409 | 51,1733193 | 7,770181555 | 20,59052131 | 11,12065246 |
| regulation of RNA biosynthetic process | 0 | 0 | 4,090437247 | 38,7977861 | 43,88266852 | 26,73113201 | 38,86213404 | 8,297292761 | 16,86306477 | 8,559619501 |
| regulation of response to stimulus | 0 | 0,602060432 | 31,82389155 | 13,94266095 | 38,19589324 | 28,49142787 | 48,53226494 | 11,95495802 | 29,32903776 | 20,93162389 |
| cell development | 0 | 3,128907002 | 22,68696283 | 0 | 37,7507475 | 14,70186049 | 45,97135412 | 13,74756888 | 18,18343963 | 18,01191993 |
| animal organ development | 0 | 1,160064586 | 25,12079478 | 0 | 40,64074983 | 19,90866261 | 62,32175922 | 15,74516087 | 21,85213484 | 16,01183396 |
| regulation of multicellular organismal process | 0 | 0 | 33,81291786 | 0 | 29,38379033 | 10,9150592 | 49,52294101 | 15,31817571 | 22,20027612 | 14,79381836 |
| positive regulation of RNA metabolic process | 0 | 0 | 3,967201441 | 23,76516538 | 31,37137346 | 24,70391119 | 34,65811405 | 6,951804466 | 17,75234153 | 9,182073821 |
| regulation of cellular process | 0 | 0 | 31,18883071 | 37,86250494 | 64,50737377 | 41,64323643 | 66,86316076 | 16,2436272 | 27,45739941 | 22,38020465 |
| macromolecule modification | 0 | 0 | 4,45920029 | 56,41753971 | 35,51924937 | 45,69205723 | 17,44358638 | 6,107525971 | 9,586557714 | 6,43328509 |
| primary metabolic process | 0 | 0 | 0 | 74,25842552 | 20,11078912 | 57,08788041 | 6,938689462 | 0,970974635 | 3,569315658 | 0 |
| positive regulation of nucleobase-containing compound metabolic process | 0 | 0 | 5,140121955 | 34,74085465 | 31,0012307 | 30,86036003 | 33,39351512 | 7,605943851 | 19,0493043 | 11,1578698 |
| cellular component organization or biogenesis | 0 | 0 | 14,63792722 | 72,98898656 | 29,41249098 | 58,43150827 | 22,70816582 | 9,780327383 | 8,108165471 | 8,208517581 |
| neurogenesis | 0,137019703 | 0,896694441 | 16,2714896 | 0 | 45,77003387 | 15,22042279 | 50,1223209 | 6,100409742 | 16,94739111 | 10,42183076 |
| regulation of biological process | 0,140698087 | 0,578904655 | 35,25598706 | 37,89662481 | 67,12088575 | 41,72286248 | 73,32100236 | 18,02408258 | 31,14727137 | 23,67132336 |
| cellular developmental process | 0,150663669 | 1,644879442 | 26,5667943 | 0,367892421 | 48,57468869 | 19,16843718 | 71,75835608 | 26,91702228 | 32,93891637 | 21,49059028 |
| cell differentiation | 0,154792434 | 1,651454021 | 26,59228736 | 0,297043057 | 48,60680871 | 19,19616378 | 71,80482439 | 26,93943925 | 32,96852862 | 21,30602303 |
| regulation of developmental process | 0,248460554 | 0 | 17,12882059 | 0,14644691 | 49,66613784 | 8,87345545 | 58,40189006 | 14,72621994 | 19,19227715 | 7,773924871 |
| positive regulation of metabolic process | 0,404756721 | 0,453633871 | 12,92103828 | 33,85926407 | 36,53804536 | 46,84415589 | 45,68835056 | 12,28695185 | 31,20497005 | 20,16305613 |
| negative regulation of cellular process | 0,554335487 | 0 | 26,42376483 | 27,01108481 | 61,31174491 | 37,93508777 | 58,89902709 | 19,38610494 | 31,83181556 | 18,91318422 |
| regulation of RNA metabolic process | 0,603066504 | 0 | 5,313566558 | 45,2807131 | 43,57049185 | 33,09189638 | 38,48203833 | 9,657683058 | 18,57700583 | 9,970971802 |
| regulation of nucleobase-containing compound metabolic process | 0,646075022 | 0 | 7,104246645 | 61,44079538 | 44,53289057 | 37,56670779 | 37,97565623 | 10,75618459 | 19,51446803 | 11,39975129 |
| regulation of macromolecule biosynthetic process | 0,682501852 | 0 | 6,009680893 | 32,56577032 | 31,84087675 | 28,79809591 | 35,2906771 | 12,7113782 | 18,43664206 | 12,05808111 |
| biological regulation | 0,79200037 | 1,167108759 | 40,35136696 | 42,40043562 | 66,62874615 | 44,62042465 | 78,35957703 | 20,70192951 | 33,79969072 | 27,4935072 |
| negative regulation of biological process | 0,799417165 | 0,473886052 | 30,75643271 | 25,29168964 | 60,38967795 | 36,53909667 | 65,41184356 | 19,19403368 | 30,71179711 | 20,3698942 |
| anatomical structure morphogenesis | 0,86404372 | 3,229455854 | 28,07671154 | 1,346493926 | 53,59652095 | 14,27147721 | 78,45729456 | 21,18929945 | 29,3222069 | 17,93030592 |
| regulation of biosynthetic process | 0,961490943 | 0 | 6,749199034 | 35,13898366 | 31,58687511 | 30,81402434 | 37,21401848 | 13,06711347 | 20,30517075 | 13,47624536 |
| positive regulation of cellular process | 0,996069973 | 2,724219874 | 31,18273953 | 39,65769463 | 64,46587862 | 63,85438618 | 86,53939416 | 25,16624358 | 48,62875955 | 33,48529041 |
| regulation of gene expression | 1,042827801 | 0 | 6,704228225 | 28,95354398 | 28,9991424 | 26,24404237 | 36,01648709 | 12,19729602 | 17,61685325 | 12,56535799 |
| regulation of macromolecule metabolic process | 1,345057605 | 0 | 12,38576066 | 50,03377835 | 41,89053405 | 41,48280996 | 37,11544172 | 13,02246106 | 21,8314559 | 15,13304768 |
| system development | 1,368005137 | 1,977431141 | 44,8316971 | 4,384624951 | 61,9560694 | 31,44436235 | 101,1854027 | 25,50951515 | 35,27683564 | 25,23504802 |
| protein localization | 1,674613853 | 0 | 8,713461167 | 43,79720079 | 18,98967787 | 55,6400242 | 11,39408774 | 5,678853941 | 8,047693917 | 7,629701067 |
| nervous system development | 1,955687444 | 1,166062795 | 23,64225584 | 2,564799848 | 51,47482531 | 22,32661936 | 63,96592667 | 12,84688956 | 19,64988705 | 12,13386762 |
| positive regulation of biological process | 2,048717862 | 3,199044712 | 32,36765952 | 37,41441508 | 62,64825847 | 64,57596019 | 90,63125729 | 25,61998169 | 49,26143175 | 35,02271929 |
| regulation of metabolic process | 2,078195297 | 0,231602808 | 15,2875089 | 60,4424418 | 43,92364223 | 45,3870895 | 45,98407079 | 16,67562484 | 26,91748366 | 20,86006892 |
| cellular macromolecule localization | 2,081350123 | 0 | 8,720248959 | 44,32434701 | 18,71881113 | 55,75529937 | 11,7986653 | 5,940327795 | 8,039315679 | 8,114110935 |
| anatomical structure development | 2,167248467 | 3,492265458 | 44,84326475 | 6,810148641 | 59,54418536 | 39,67436708 | 100,8083892 | 31,863002 | 44,13073631 | 36,13566377 |
| multicellular organism development | 2,41324671 | 2,568301247 | 43,50186491 | 8,297317254 | 61,54165508 | 37,76236419 | 102,6454279 | 28,78442797 | 38,58507328 | 27,11730767 |
| regulation of cellular component organization | 2,443584755 | 0 | 22,06457973 | 22,19671709 | 35,4971857 | 39,11955166 | 31,45033919 | 9,193000856 | 13,08511168 | 14,53361463 |
| developmental process | 2,490271094 | 3,091809428 | 46,1061144 | 9,243379714 | 60,30667805 | 41,88992644 | 97,56591867 | 35,43533229 | 46,65158542 | 34,77306073 |
| macromolecule localization | 2,571292501 | 0,339459474 | 11,62760453 | 41,44182675 | 19,25808064 | 61,06591055 | 14,06183157 | 7,0508912 | 12,13413427 | 10,8281941 |
| transport | 2,747595836 | 2,313266038 | 23,77933892 | 16,57385839 | 23,51001549 | 40,81027925 | 36,13114774 | 17,91520733 | 30,50751172 | 21,57777657 |
| cellular localization | 3,122086677 | 2,169941242 | 12,46083882 | 51,62587996 | 21,83985699 | 61,8177776 | 24,39708699 | 10,95395897 | 15,07420202 | 12,25768768 |
| regulation of primary metabolic process | 3,290654914 | 0,105747463 | 17,03820221 | 74,81456444 | 57,69411391 | 55,60300391 | 52,16444055 | 13,94377533 | 29,30668301 | 18,03356275 |
| organelle organization | 3,480372612 | 0,684015209 | 8,324556673 | 87,40589811 | 28,33575228 | 69,06186536 | 15,19724154 | 11,71882731 | 5,586584915 | 21,71660631 |
| establishment of localization | 3,554786079 | 2,064759283 | 25,5835924 | 21,63774338 | 24,99564133 | 49,50294952 | 34,93514884 | 17,82552997 | 29,43279674 | 22,11787276 |
| protein metabolic process | 3,778537658 | 0 | 8,467088063 | 65,57236363 | 34,16922589 | 64,40631311 | 21,25908579 | 8,023811093 | 13,50392015 | 15,01170554 |
| localization | 4,588505543 | 5,10332045 | 29,49530017 | 34,11369608 | 32,22225975 | 63,4181308 | 44,28117229 | 20,06061893 | 33,72955706 | 28,94318831 |
